# Macrophages use sub-micron actin ridges in phagocytosis of encapsulated fungal pathogens

**DOI:** 10.64898/2026.10.30.763038

**Authors:** Jaime Cañedo, Mahrukh Shameem, Laura Martinez-Campesino, Chiara Niespolo, Taghreed Shamrani, James Bradford, Yin Xin Ho, Lynne R. Prince, Jason S. King, Endre Kiss-Toth, Heather L. Wilson, Rhoda J. Hawkins, Simon A. Johnston

## Abstract

Ingestion and killing of pathogens by phagocytic cells is a key aspect of animal immunity. *Cryptococcus neoformans* is a major pathogen of AIDS patients that has a polysaccharide capsule that is critical for virulence. Phagocytic function is necessary for normal immunity to cryptococcal infection, and we sought to determine how *Cryptococcus* capsule prevents phagocytosis. In doing so, we discovered a new sub-micron actin structure, the ridge, that is present on macrophages, neutrophils and amoebae. Macrophages preferentially bind cryptococci via ridges and suppression of ridges is sufficient to prevent phagocytosis. Ridges are present on macrophages *in vivo* and are lost in interleukin 10 stimulated macrophages. The discovery and functional description of macrophage surface ridges fills a critical gap in our understanding of how phagocytes ingest encapsulated fungal pathogens, a crucial step in treating infection.

## Introduction

Professional phagocytes are one of the earliest and most well-conserved eukaryotic cell types. They exist today as free-living amoebae and phagocytic immune cells present in all animal lineages (1). *Cryptococcus neoformans* is an environmental yeast and that can cause fatal meningoencephalitis in immunocompromised humans. Susceptibility to fatal disease is characterised by an inability of immune cell macrophages to phagocytose and kill cryptococcal cells (2-5).

A universal characteristic of phagocytic cells is their highly dynamic and varied actin cytoskeletal structure. Actin dynamics are critical to macrophage function: cell migration to a target, its internalisation through phagocytosis, extravasation and extracellular matrix degradation. Phagocytosis requires deformation of the phagocytic cell to enwrap and ingest their target into a new unilamellar compartment, the phagosome. Many pathogenic microbes possess a polysaccharide capsule that has been implicated in the inhibition of this process. Cryptococcus is unique among the human fungal pathogens in being encapsulated. We and others have demonstrated how the cryptococcal capsule might prevent phagocytosis but the interaction of macrophages with cryptococci has not be sufficiently resolved to define the interacting surface (5-7). Therefore, we performed ultrastructural scanning electron microscopy and beyond-diffraction limit fluorescence microscopy of the highly delicate capsule and macrophage membrane that allowed measurement with high precision of the interactive surface of cryptococci with macrophages during phagocytosis.

## Results

### Macrophage sub-micron ridges are the primary interactive surface with cryptococci

High resolution imaging of the macrophage surface revealed that sub-micron apical surface protrusions were the dominant feature of the macrophage *Cryptococcus* interaction (Fig 1A-G, Table S1). This was either by sheet-like ridges or finger like protrusions (filopodia), but ridges were vastly more frequent than filopodia (Fig. 1H, Table S2). Antibody opsonisation increased the phagocytosis of cryptococci by macrophages (Fig 1L) (8). We hypothesised that this would remove the requirement for ridges in the binding of cryptococci by ridges as phagocytic Fc receptors could directly bind the capsular surface via capsule reactive antibody, removing the requirement for cell wall binding pattern recognition receptors. In support of this hypothesis the increase in phagocytosis could be explained by a three-fold increase in binding via protrusion-less macrophage surfaces (Fig. 1L, Table S3), while sub-micron protrusions remained the dominant interacting feature. Thus, we identified three common actin-based membrane protrusions used by macrophages in their interaction with *Cryptococcus*: filopodia (Fig. 1A and 1D), surface ridges (Fig. 1B and 1E) and dorsal ruffles (Fig. 1C and 1F). Though surface ridges and dorsal ruffles are both sheet-like structures, their morphologies could be easily differentiated using SEM as ruffles are larger with a more complex architecture than ridges (Fig. S1E green arrow and S1F green box, Table S5).

**Figure 1.**
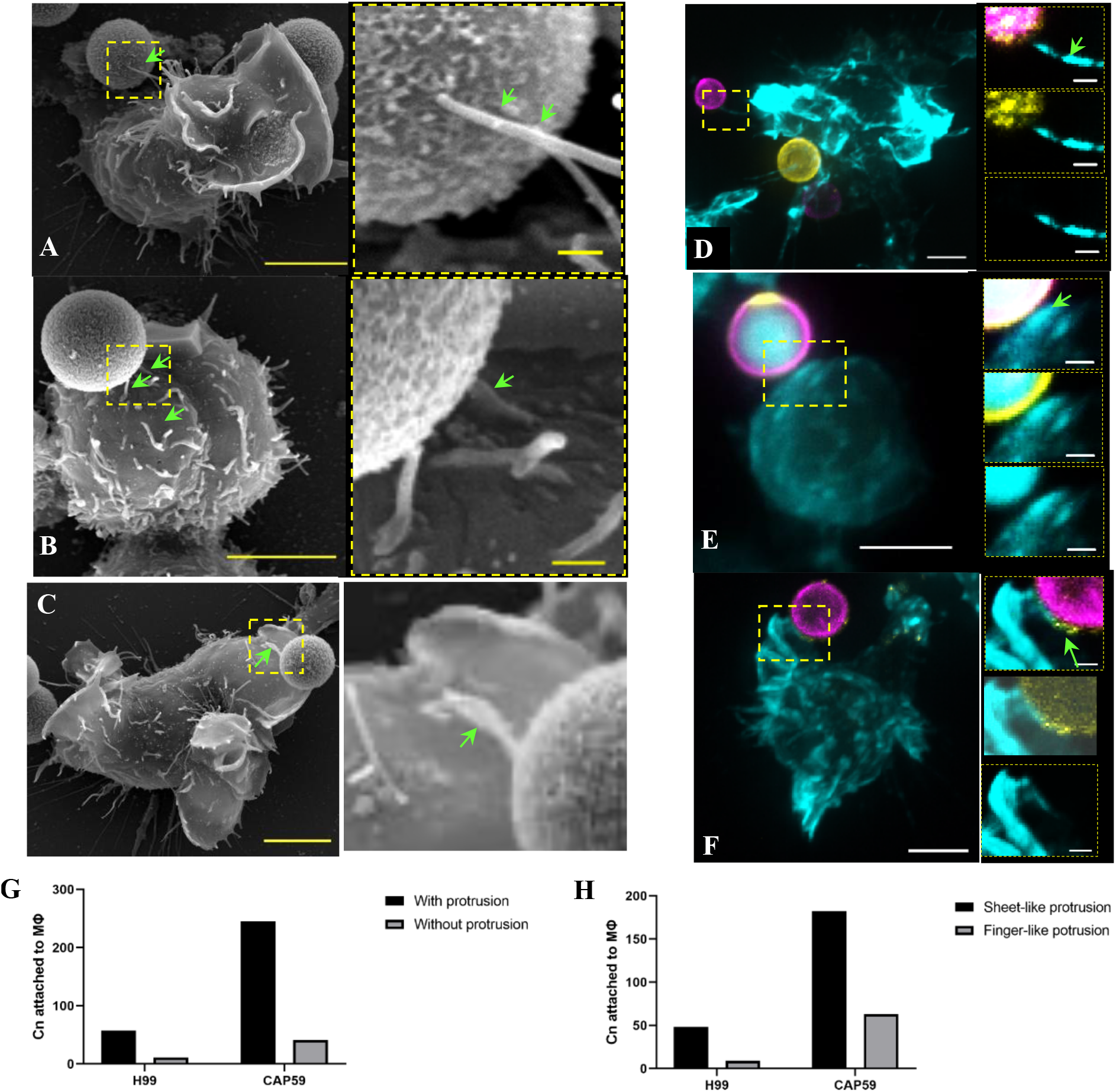
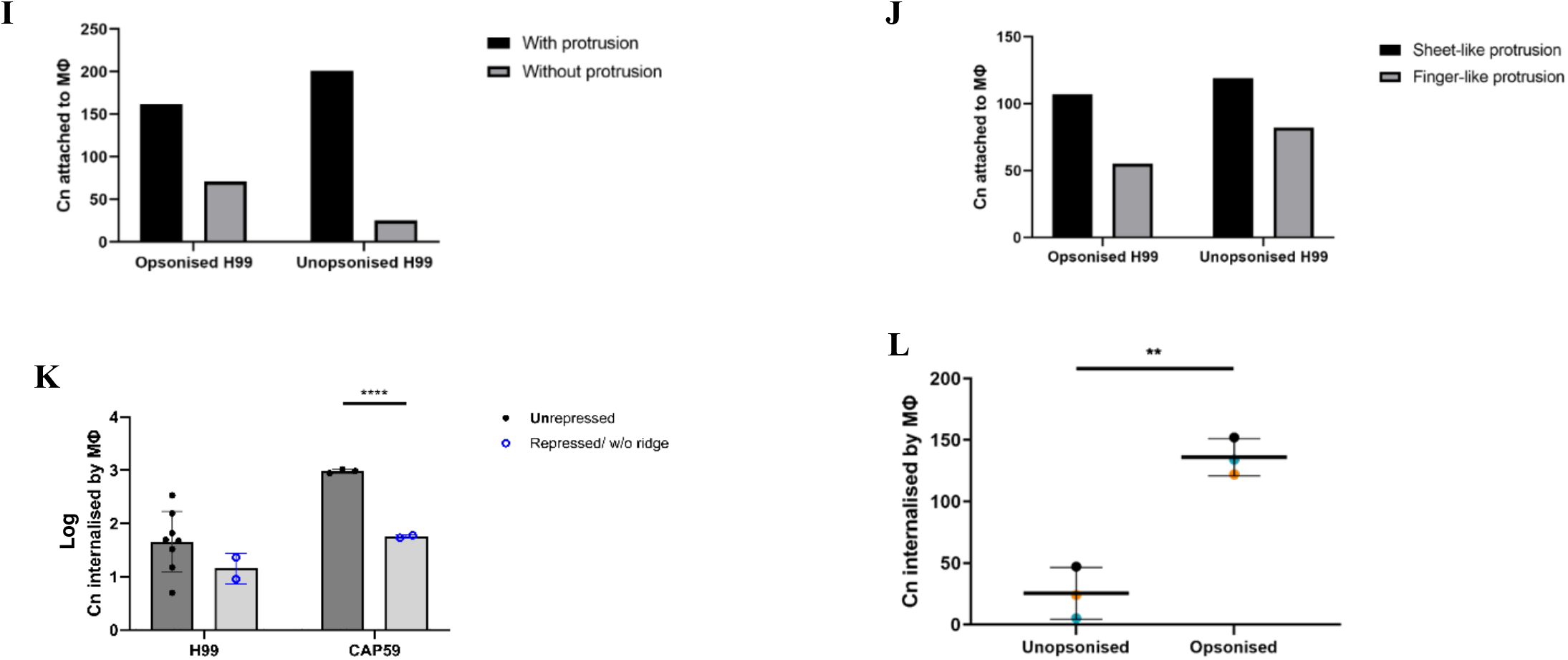
Surface ridges are the interaction surface of macrophages with cryptococci. **A-C**, Ultrastructural features of macrophage and *Cryptococcus neoformans* (Cn) interaction *in vitro* through scanning electron microscopy: **(C)** J774 macrophage contacting a cryptococcal cell by filopodium (green arrow), **(B)** by surface ridge (green arrow), **(C)** by membrane ruffle (green arrow), **D-F**, deconvolved widefield fluorescence images of macrophage and *Cryptococcus* interaction *in vitro* (cyan – f-actin, yellow – capsule, magenta – cell wall): **(D)** showing contact with actin filopodium (green arrow), **(E)** with actin surface ridge (green arrow), **(F)** with actin ruffle (green arrow), (**G)**, *Cryptococcus* showed higher attachment to macrophage surface with actin protrusion than without actin protrusion independent of capsule possession (representative numerical data from n = 3 (>500 macrophages scored per repeat), dependence of relationship on possession of capsule *p* = 0.704 two-sided Fisher’s exact test (contingency table in supplementary table S1), **(H)**, *Cryptococcus* had higher attachment to sheet-like protrusion than finger-like actin protrusion of macrophages (representative numerical data from n = 3 (>500 macrophages scored per repeat) regardless of capsule possession (*p* = 0.124) as shown by two-sided Fisher’s exact test (contingency table in supplementary table S2), **(I)**, opsonised Cryptococci had increased attachment to protrusion-less macrophage surface (*p* < 0.0001) as confirmed by two-sided Fisher’s exact test (contingency table in supplementary table S3), **(J)**, both unopsonised and opsonised H99 Cryptococci were also found to have higher attachment to actin protrusion with sheet-like geometry than finger-like ones (*p* = 0.124) as revealed by two-sided Fisher’s exact test (contingency table in supplementary table S4) , **(K)**, loss of surface ridges impaired uptake of acapsular Cryptococcus (CAP59) by macrophages as revealed by multiple unpaired t tests (*t*(3) = 37.52, \*\*\*\**p* < 0.0001, *n* = 2-8 repeats (>500 macrophages scored per repeat), bars: *SD*), **(K)**, Opsonisation of H99 Cryptococcal cells increased its internalisation by MΦ as confirmed by unpaired t test (*t*(4) = 7.403, \*\**p* = 0.002), *n* = 3 (each colour represents an experimental repeat), bars: mean (*M*) ± standard deviation (*SD*). Scale bars, 5 μm **(A-E** main panel); 500 nm **(A-C** zoomed-in panel/yellow box); 1 μm **(D**,**E** zoomed-in panel).

### Loss of macrophage ridges is sufficient to inhibit phagocytosis of cryptococci

As early as 30 minutes post exposure with cryptococci we observed cytoskeletal remodelling of macrophages dominated by large macropinocytic ruffles (Fig. S1B, S1E arrows, Fig. S1L yellow arrows). By two hours, neither ridges nor ruffles were present (Fig. S1C, Fig. S1M), changes that did not occur when challenging cells with similar sized latex beads (Fig. S1D, S1M). We also looked at the events within 30 minutes of infection as the macrophage surface seemed to transition from a ridge-enriched to a dorsal ruffle-enriched region. Using single cell SEM, we profiled the density of surface ridges on macrophages during infection using an annotated image analysis pipeline (Fig. S2B, S2E; (9)) whilst manually annotating and isolating the dorsal ruffles (Fig. S1E and Fig. S1F) to determine their area densities. Strikingly, our results confirmed that decrease in surface ridges during infection was concomitant with increase in dorsal ruffles in both capsular (Fig. S1G) and acapsular cryptococcal infection (Fig. S1H). This also revealed that the peak density of ruffles on the surface at ∼20% (Fig. S1G and S1H purple line, 30 minutes) was very similar to the peak density of ridges (Fig. S1G and S1H orange line, 0 minute), and the peak ruffle expression at 30 minutes coincided very well with the lowest ridge density (Fig. S1G and S1H, compare purple and orange lines at 30 minutes), suggesting conservation of actin in producing cellular protrusions during infection. Therefore, to determine if ridges were a requirement for successful phagocytosis of cryptococci, we measured phagocytosis in the absence of ridges 30 minutes post-exposure to cryptococci. We found that uptake was reduced for both wild-type capsular H99 cryptococci and derived acapsular cap59 mutant (Fig. 1K).

### Actin ridges are found on mammalian professional phagocytes and amoebae

We next profiled the density and nanoscopic geometry of surface ridges across professional phagocytes. We found that in both primary and cell line phagocytes cultured *in vitro*, neutrophils and macrophages, and amoebae (Fig. 2E – 2L), ridge area density varied across cell type (Fig. 2O): human neutrophils and mouse bone marrow-derived macrophages showed the highest ridge density, constituting about 20% of their total cell transverse cross-sectional area, whilst the microbivore *Dictyostelium discoideum* amoeba had the lowest. Strikingly, undifferentiated pre-monocytic THP1 cells (Fig. 2K) did not possess ridges but they were present post-differentiation to macrophage-like cells (Fig. 1L), further implicating ridges to a phagocytic lifestyle. In all professional phagocytes we examined, the thickness of surface ridges (Fig. S2P – S2S) were very consistent and peaked at around 140 nm.

**Figure 2.**
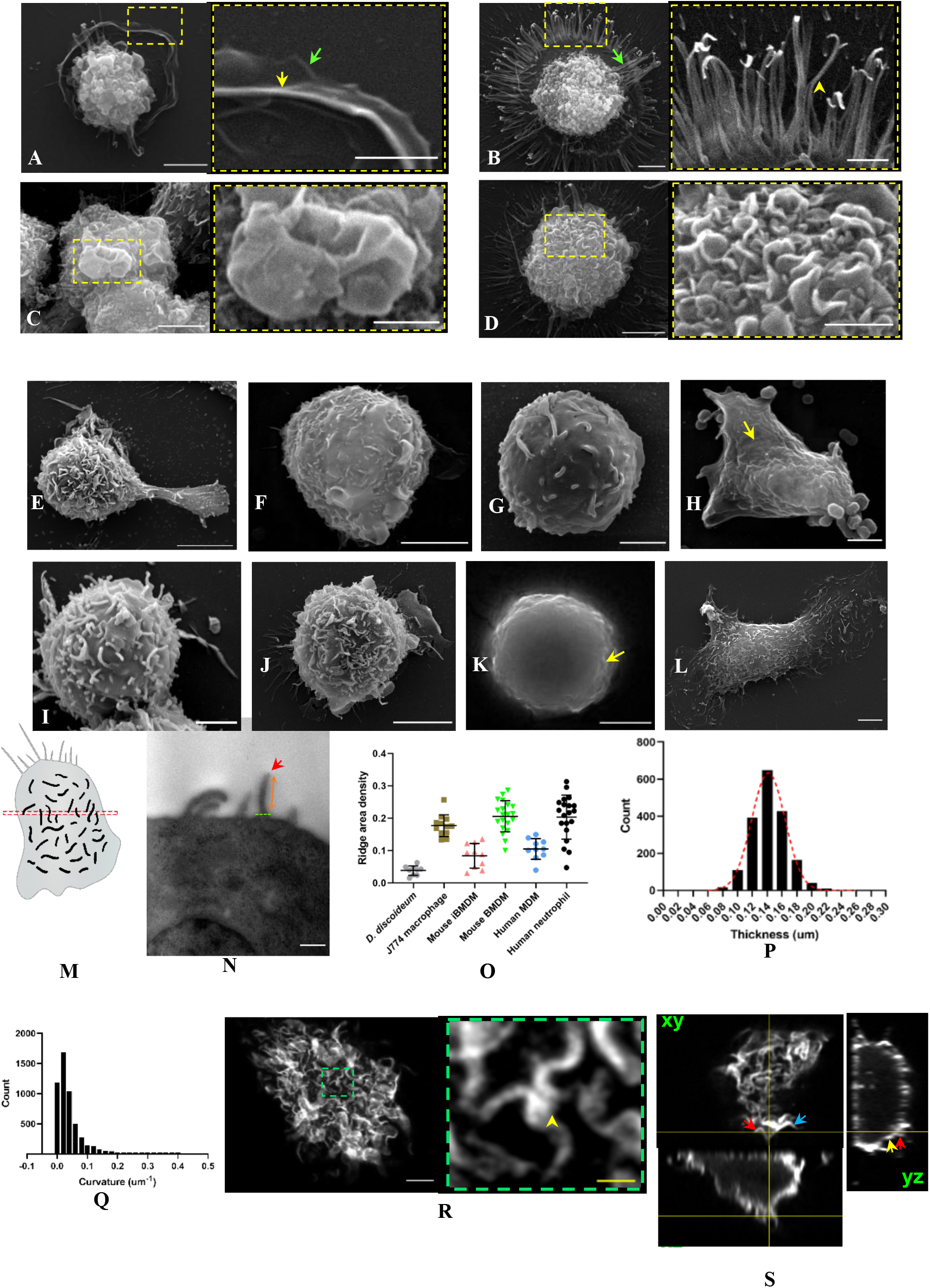
Surface ridges are a highly conserved actin-rich sheet-like protrusion observed *in vitro* and *in vivo* across different phagocytes. **a-d**, Scanning electron microscopy (SEM) of different types of surface protrusions produced *in vitro* by unstimulated human monocyte-derived macrophage (HMDM) and its zoomed-in section shown in the adjacent yellow box: (**a)** *pseudopodia* showing its planar and sheet-like features with a secondary dorsal protrusion (yellow arrow) close to its border (green arrow), **(b)** *filopodia* emerging as finger-like protrusions from the cell body, **(c)** *membrane ruffles* as sheet-like protrusions and forming a macropinocytic cup, **(d)** *surface ridges* showing dense and sheet-like features on dorsal cell surface, **e-l**, SEM of cell surfaces of different types of phagocytes cultured *in vitro*: **(e)** mouse bone marrow-derived macrophage (BMDM), **(f)** immortalised mouse (BMDM), **(g**) human neutrophil, **(h)** *Dictyoselium discoideum* (NC4), **(i)** RAW264.7 macrophage-like cell, **(j)** J774 macrophage-like cell, **(k)** undifferentiated THP-1 cell, **(l)** differentiated THP-1, **m**, schematic diagram of a phagocyte cross-section to illustrate ridge architecture examined by transmission electron microscopy, TEM, (top - top view of a macrophage surface showing a thin cross-sectional region (red box) used for TEM examination, bottom – a phagocyte’s cross-sectional body diagram (red box in top image), **n**, TEM image of the apical surface of a J774 cell cultured *in vitro* showing the cross-sectional features of a surface ridge (green - base, orange -body, red - tip), **o**, ridge area density of different phagocytes computed through an image analysis routine, *ridge detection* (RD), of SEM images (supplementary figure S2b) showed statistically significant differences between cell types as revealed by ordinary one-way ANOVA (*F(5*,*75) = 26*.*99, p < 0*.*0001, n* = 9-20 cells scored per cell type, bars: mean (*M*) ± *SD*), for Tukey’s test comparison between groups, refer to supplementary table 1.1, **p**, combined ridge thickness distribution of all J774 cells examined (*n =* 12 cells (1818 ridges), *M* = 0.142, *SD* = 0.023) with Gaussian fit (R^2^ = 0.997) using SEM images and RD analysis, **q**, ridge bending modulus using J774 cells (*n* = 58 ridges (5220 curvature points), *M* = 0.04, median = 0.02) derived by SEM and a custom-built script (supplementary figure g-i), **r**, airyscan superresolution laser scanning imaging of f-actin in the apical region (about 1 μm z thickness) of J774 cell using phalloidin-TRITC probe reveals the actin component of surface ridges, zoomed in panel (green box) shows the bendy morphology of actin in these protrusions, **s**, orthogonal examination of f-actin cytoskeleton of a J774 cell reveals surface ridges (red and blue arrows) emanate from and continuous with the actin cortex (yellow arrow). Scale bars, 5 μm **(a-d, e, f, j, k, l, r** main panel); 2 μm **(a-d** zoomed-in panel/yellow box, **g, h, l)**; 1 μm **(r** zoomed in panel/green box); 500 nm **(n)**.

### Actin ridges have a geometry indistinguishable to other macrophage surface actin-based structures

As noted above, phagocytes exhibit diverse membrane protrusions supported by an inner actin scaffold (Fig S4A – S4J; (10-13) . We were struck by the apparent similarity in thickness between the different macrophage surface actin-based structures: lamellipodia (Fig. 2A), filopodia (Fig. 2B) and ridges (Fig. 2D). To confirm this consistent actin protrusion thickness, we next undertook a comparative analysis. Lamellipodia (Fig. 2A and Fig. S4D – S4G) were seen as flat, sheet-like protrusions extending from the phagocyte’s body (Fig. S4D,G green arrows) (14,15). We manually defined lamellipodial protrusions from SEM (Fig. 2A yellow arrow) and fluorescence images (Fig. S4G green arrows), and used the ridge detection-based (9) image analysis pipeline we previously implemented (Fig S2B) to determine their thickness and area densities. Similarly, we used the same pipeline to analyse the actin thickness of filopodia (Fig. S4C), surface ridges (Fig. S4B green arrow) and cortical actin (Fig. S4B red arrow) in macrophages. Secondary manual segmentation from SEM (Fig. S4O) and actin labelled fluorescent images (Fig. S4P), found no difference in the thicknesses of lamellipodia, filopodia, ridges, and the actin component of the cortex, consistent with recent measurements (16,17). Like other macrophage surface actin features, ridges were highly dynamic with high turnover on the cell surface, mean lifetime of 55 seconds (Fig. S5V), slight curvature (Fig 2Q; Fig. S2D; Fig. S2I), splitting (Fig. S5G – S5N) and fusing (Fig. S5A – S5F) behaviours whilst remarkably maintaining their thickness (Fig. S5S, S5U, S5O – S5Q).

### Surface ridges are repressed by IL-10 and tissue environment

As macrophages are sensitive to inflammatory factors, which highlights their critical role in immune response, we tested how macrophage ridges were regulated by different types of inflammatory mediators and by local tissue environment by SEM imaging of resident macrophages. We stimulated human monocyte-derived macrophages with different pro-inflammatory (LPS/IFNγ) and anti-inflammatory (IL-4 and IL-10) cytokines, and measured changes in cell surface area, aspect ratio (elongation) and ridge density. Our results showed that cell surface area was maintained in all activation conditions (Fig. 3E), whilst ridge density was found to be more variable (Fig. 3F) suggesting independent mechanisms that control cell size and surface ridge production. None of the cytokines further increased ridge density which indicates a limit on ridge production that is potentially due to limited membrane reservoir in macrophages. Notably, IL-10 almost completely inhibited ridge production, and transformed the macrophage surface into a wrinkled surface phenotype (Fig. 3D) observed previously in mast cells (18).

**Figure 3.**
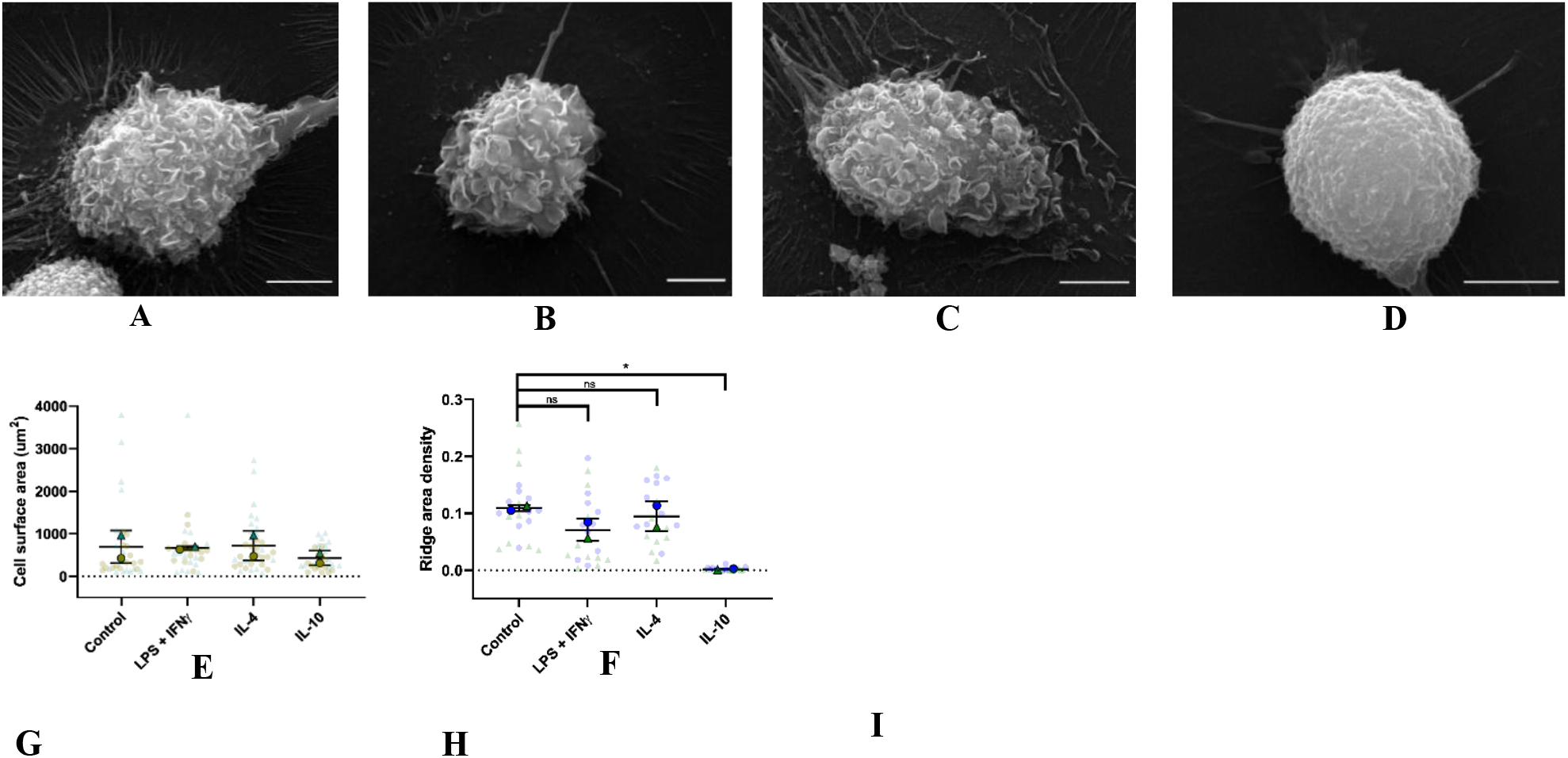
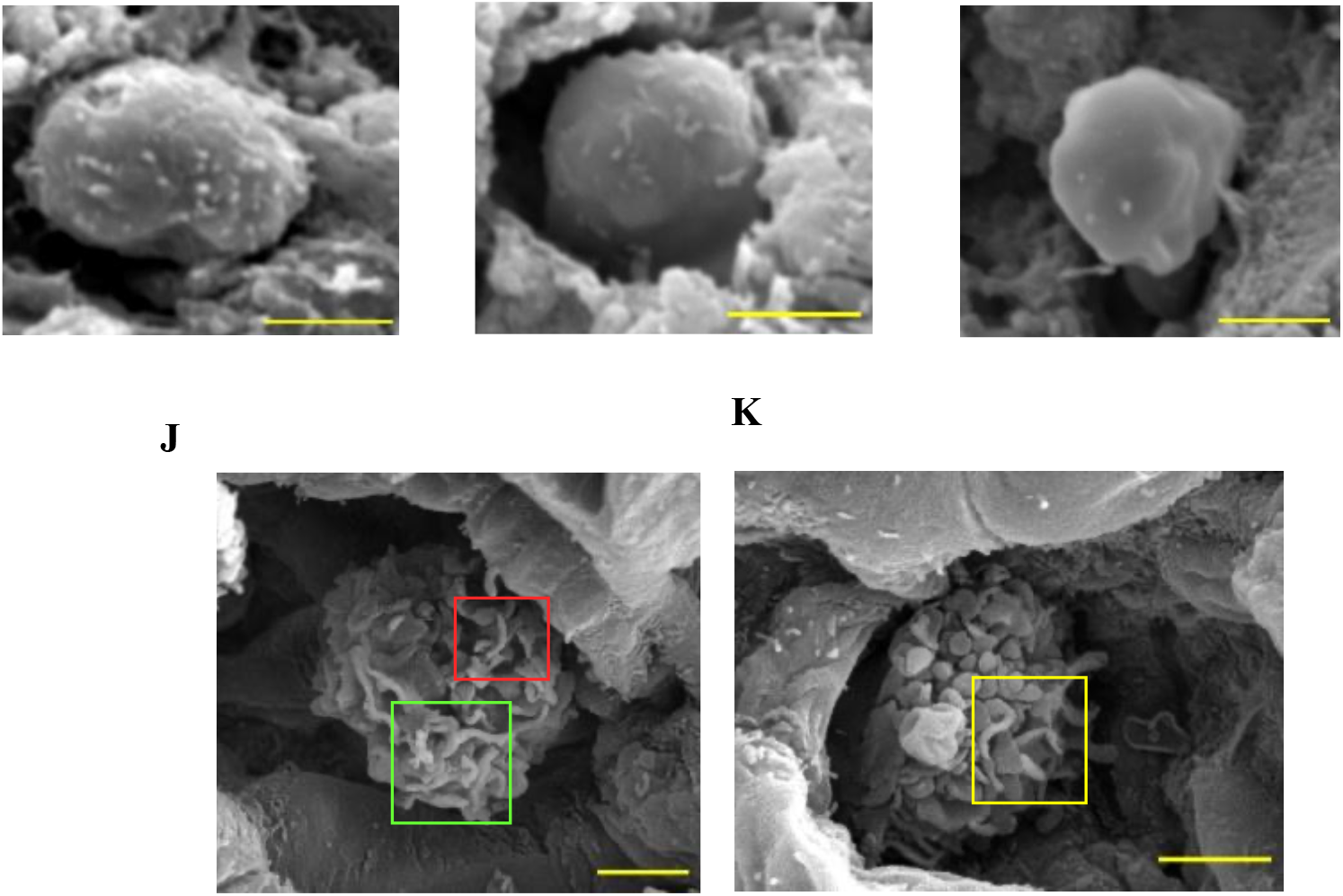
Surface ridges are inhibited by IL-10 cytokine and by suppression of pro-inflammatory genes. **a-d**, SEM of HMDM surfaces *in vitro* at different activation states: **(a)** unstimulated/control, **(b)** activated by LPS/IFNγ, **(c)** by IL-4, **(d)** by IL-10 showing almost complete inhibition of surface ridges **e**, no statistically significant change in cell surface area of HMDM was found following activation with different cytokines as confirmed by ordinary one-way ANOVA (*F*(3,4) = 0.469, *p* = 0.72), *n* = 2 (14-17 cells per repeat), bars: *M* ± *SD*), **f**, surface ridges of HMDM were completely inhibited upon *in vitro* induction with IL-10 cytokine as shown by ordinary one way ANOVA (*F*(5,4) = 16.04, *p* = 0.011), *n* = 2 (10 cells per repeat), bars: *M* ± *SD*), (**G-I)**, surface features of mouse liver resident macrophage, Kupffer cell, viewed in SEM *in vivo*, (**J**,**K)**, surface features of mouse alveolar macrophage *in vivo* revealing the different types of protrusion they produce in their natural environment (green – surface ridges, red – mushroom-shaped protrusions, yellow – membrane ruffles), Scale bars, 5 μm **(A-D)**; 2 μm **(G-K)**.

### Ridges exist in vivo and maintain IL 10 sensitivity

Because IL-10 is a well-characterised cytokine widely expressed in the liver (19, 20), we then examined the resident liver macrophages, Kupffer cells, in mice *in vivo*. We found their surface rarely possessed ridges, supporting our finding of an inhibitory property of IL-10 to surface ridges (Fig. 3G-I). Finally, we hypothesised that alveolar macrophage, given their sentinel role in the lungs (21, 22) would be a likely cell type to confirm their structure and function *in vivo*. As for Kupffer cells we were able to use perfusion fixation to preserve alveolar macrophage *in vivo* cell surface feature for examination *ex vivo*. Mouse alveolar macrophages were indeed dominated by surface ridges *in vivo*, with remarkable correspondence to our findings *in vitro* and demonstrating their potential *in vivo* function in phagocytosis as we have shown *in vitro* (Fig. 1).

## Discussion

The ability of professional phagocytes to rapidly and effectively internalize microbes is fundamental to animal immunity. Our findings define a previously undescribed, sub-micron actin-based membrane protrusion, the surface ridge, which is critical for the phagocytosis of the encapsulated fungal pathogen *Cryptococcus neoformans*.

Our ultrastructural and super-resolution analysis demonstrates the macrophage ridges as the principal feature facilitating the initial interaction between macrophages and *C. neoformans*, whether opsonised or unopsonised. The ridge’s sheet-like geometry provides an effective mechanism to engage the cell wall despite the overlying capsule, acting as a crucial intermediate structure to initiate phagocytosis. This mechanism likely overcomes the low-affinity, distributed binding of phagocytic receptors, a task that larger, flatter membrane surfaces fail to achieve effectively against an encapsulated target. Beyond their specific role in fungal recognition, we establish the surface ridge as a highly conserved and ubiquitous feature across diverse professional phagocytes, including human and mouse macrophages, neutrophils, and even the microbivore amoeba *Dictyostelium discoideum*. This conservation across vast evolutionary distance suggests a fundamental and ancient role for this geometry in phagocytic processes. Furthermore, the remarkable consistency in the ridge thickness across all examined cell types, matching that of other core actin structures, cortical actin, lamellipodia, and filopodia, points to a shared, tightly regulated biophysical mechanism governing the polymerization and stability of these membrane structures. Like other actin structures, we found that ridges are highly dynamic, capable of splitting and fusing while maintaining their invariant thickness, highlighting a sophisticated, self-organizing actin machinery.

Crucially, our study places the surface ridge within the broader context of immune regulation and pathogenesis. We demonstrate an inverse relationship between surface ridge density and the expression of macropinocytic ruffles during active infection. Intriguingly, this change may reflect a pathogen-driven diversion of the actin machinery from ridge mediated phagocytosis toward fluid-phase macropinocytosis which *C. neoformans* may subvert for its own benefit, consistent with the known dichotomy of macrophage outcomes in cryptococcal disease and the evolution of *C. neoformans* virulence due environmental phagocyte interactions.

Finally, we identify a direct link between immune polarization and ridge expression. The anti-inflammatory cytokine IL-10 nearly abolishes ridge production, converting the macrophage to a smooth, wrinkled surface. This *in vitro* observation was strongly supported by our *in vivo* profiling, where Kupffer cells, resident liver macrophages known to operate in an IL-10-rich, anti-inflammatory environment, exhibit a distinct lack of surface ridges. Conversely, highly active alveolar macrophages, constantly exposed to environmental challenges, are dominated by these protrusions, with implications for tissue-specific susceptibility to infection.

In summary, the identification and characterization of the macrophage surface ridge not only fills a critical gap in our understanding of how phagocytes manage difficult-to-internalize pathogens but also uncovers a novel, evolutionarily conserved, and inflammation-sensitive actin structure. Future work should focus on identifying the specific molecular machinery that regulate the formation, dynamics, and IL-10-mediated repression of the surface ridge, thus opening new avenues for therapeutic intervention in opportunistic fungal infections and the manipulation of phagocytic processes.

## Materials and Methods

### Cell types

The following cell types were used in this project: J774, RAW264.7, THP1, human monocyte derived macrophages (MDM), human neutrophils, mouse bone marrow-derived macrophages (BMDM), immortalised BMDM, mouse alveolar macrophages and *Dictyostelium discoideum*.

J774 (23) and RAW264.7 (24) macrophage-like cell lines were recovered from cryovial stocks stored in liquid nitrogen and thawed for ∼2 min in a 37 °C water bath. Cells were resuspended in 4 ml complete DMEM: 88% DMEM (Sigma D5546), 10% fetal bovine serum (FBS) (Sigma F7524), 1% 2 mM L-glutamine (Sigma G7513) and 1% penicillin–streptomycin (100 U ml^−1^ penicillin G and 0.1 mg ml^−1^ streptomycin; Sigma P4333), centrifuged for 5 min at 1000 × g, and resuspended in ≥10 ml complete DMEM. Cells were cultured at 37 °C, 5% CO_2_, passaged at ≥80% confluency, and experiments were performed using cells between passages 3 and 14.

Undifferentiated THP-1 cells were maintained in complete RPMI: 89% RPMI (BioWhittaker), 10% FBS (Sigma F7524) and 1% penicillin–streptomycin (Sigma P4333). Cell density was maintained above 1 × 10^5^ cells ml^−1^ and cells were passaged up to 20. For differentiation, THP-1 cells were centrifuged at 1,250 rpm for 5 min, resuspended in fresh complete RPMI, counted and seeded onto glass coverslips in 24-well plates at 2 × 10^5^ cells per well in 500 µl complete RPMI supplemented with 50 ng ml^−1^ phorbol 12-myristate 13-acetate (PMA). Cells were incubated for 3 days at 37 °C, 5% CO_2_, followed by replacement with fresh complete RPMI and incubation for a further 2 days.

Whole blood was mixed with trisodium citrate dihydrate at a 9:1 (citrate:blood) ratio and peripheral blood mononuclear cells (PBMCs) were isolated by density centrifugation using Ficoll. Monocytes were enriched by positive selection using human anti-CD14 microbeads (Miltenyi Biotec 130-097-052). Monocytes were centrifuged at 1500 rpm for 5 min and resuspended in complete RPMI with 100 ng ml^−1^ recombinant human M-CSF (rhM-CSF), then incubated at 37 °C, 5% CO_2_ for 7 days to differentiate into macrophages. For polarization, HMDMs were stimulated overnight with 100 ng ml^−1^ LPS + 20 ng ml^−1^ IFNγ , or 20 ng ml^−1^ IL-4 or 20 ng ml^−1^ IL-10, before processing for scanning electron microscopy (SEM) as described below. For TRIB1 knockdown, differentiated monocytes were prepared for transfection as described previously (25).

Human neutrophils were isolated from whole blood from healthy volunteers by Plasma–Percoll density centrifugation as previously described (26). Freshly isolated neutrophils were resuspended at 5 × 10^6^ cells ml^−1^ in RPMI (Thermo Fisher 1640) supplemented with 10% (v/v) heat-inactivated FBS (PromoCell) and 25 mM HEPES (Sigma Aldrich). Cells were allowed to settle onto uncoated glass coverslips in 24-well plates for several hours at 37 °C, 5% CO_2_ prior to SEM processing.

Wild type (C57BL/6) and *Trib3*^*+/−*^ mice (27) or Trib1-/-mice (28) were used to examine bone marrow-derived macrophages. Briefly, mice were euthanised and femurs and tibias were collected. Cell suspensions were centrifuged at 500 × g for 5 min, resuspended in complete DMEM supplemented with 10% L929 cell-conditioned medium, seeded into 24-well plates and incubated at 37 °C, 5% CO_2_ for 5 days, after which medium was replaced prior to SEM processing.

Immortalised BMDMs (iBMDMs) were kindly provided by D. Brough (University of Manchester, generated as described (29) and cultured in complete DMEM with 10% FBS and incubation conditions as above. At ∼90% confluency, cells were passaged, detached by scraping in fresh medium and seeded into 24-well plates at 1 × 10^6^ cells per well for SEM examination.

For isolation of mouse alveolar macrophages, mice were euthanised by intraperitoneal injection of sodium pentobarbital and confirmation via exsanguination of the femoral artery. The trachea was cannulated, secured and washed twice medium consisting of 0.4% EDTA, 0.5% FBS and 99.1% PBS. Samples were centrifuged at 250 × g for 10 min at 4 °C, and the pellets were resuspended in PCLS medium (90% high-glucose DMEM, 9% FBS, 1% penicillin–streptomycin). Cells were incubated overnight at 37 °C, 5% CO_2_ before SEM processing.

*Dictyostelium discoideum* (NC4 strain) were grown with *Klebsiella aerogenes* in adherent culture at 22 °C, allowed to settle on coverslips for 30 min, fixed with 4% paraformaldehyde (PFA) for 5 min at room temperature before processing for SEM.

### Tissue samples

For the isolation of mouse lung and liver, mice were deeply anaesthetised by intraperitoneal injection of sodium pentobarbital. An incision is made to remove the ribs, exposing the heart. A blunt perfusion needle is injected into the left ventricle and the right atrium is cut. The animal is perfused with 10mL of PBS, followed by 20mL of 4% PFA. Once the entire animal was fixed and the perfusion solution was empty, the lungs and liver were removed and placed in 4% PFA for 10 min at room temperature. Tissues were washed three times in PBS, then fixed with 2.5% glutaraldehyde in 0.1 M sodium phosphate buffer overnight at 4 °C. Tissues were mounted and sectioned using a vibratome to obtain 100 µm sections, washed at least three times before processing for SEM.

### Scanning electron microscopy

Cells on uncoated glass coverslips were fixed in 4% PFA for 10 min at room temperature, washed three times with PBS, and fixed in 3% GA in 0.1 M sodium phosphate buffer overnight at 4 °C. Samples were washed twice in 0.1 M phosphate buffer, post-fixed with 2% aqueous osmium tetroxide for 1 h at room temperature, and dehydrated through graded ethanol (75%, 95%, 100% twice). Samples were critical point dried using a Leica EM CPD300 using the following settings: slow CO_2_ influx, 2 min delay prior to 24 exchange cycles, slow heating to 34 °C, and slow gas out at 75% of normal speed. Dried samples were mounted on aluminium stubs and sputter coated with ∼25 nm gold. Samples were imaged using a TESCAN Vega3 LMU scanning electron microscope at 10–30 kV accelerating voltage, 9 mm working distance, 63 nm spot size, and 0° stage tilt. Magnifications ranged from 500× to 90,000×, with typical settings of ∼500× for cell populations, 10,000–30,000× for single cells, and 30,000–90,000× for membrane protrusions.

For 3D reconstruction of macrophage dorsal surfaces, Alicona MeX v6.2 software was used for photometric stereo-based SEM reconstruction. Automated angular calibration was performed at 0°, +45° and −45° using ridge features at 10,000– 30,000×. 3D renderings were generated with automatic offset correction, and ridge geometry (width, length and height) was quantified from rendered profiles.

### SEM image processing and quantification

Ridge detection-based morphometry was done through Fiji (30). Ridge measurements were rendered using ridge detection plug-in (9) with the following parameters: sigma 1–3, lower threshold 0, upper threshold 1–6. Automated outputs were visually inspected to confirm accurate detection and manual annotations were additionally performed in TESCAN Vega3 LMU SEM software and compared with ridge detection-derived values (Fig S2E). Individual ridges were manually extracted by tracing ridge borders, after which ridge detection (9) was used to determine ridge midlines. Ridge curvature was quantified using a custom ridge curvature profiling script (code source below). Curvature values were summarised as histograms for individual ridges and pooled ridge sets.

Low magnification SEM images (∼500×) were used to quantify cell surface area and elongation (aspect ratio). Cells were segmented by intensity thresholding in Fiji, followed by manual inspection to confirm correct border identification. Particle analysis was applied to extract cell surface area and aspect ratio values.

### Transmission electron microscopy

Seeded cells on glass coverslips (1 × 10^5^ per well) were fixed in 3% GA in 0.1 M sodium phosphate buffer overnight at 4 °C, washed twice in 0.1 M phosphate buffer, post-fixed in 2% aqueous osmium tetroxide for 1 h at room temperature, and dehydrated in ethanol (75%, 95%, 100% twice). Samples were incubated in propylene oxide twice for 15 min each. Specimens were infiltrated overnight in a 50:50 propylene oxide:araldite mixture at room temperature, followed by incubation in 100% araldite for 6–8 h and embedding in fresh araldite for 48–72 h at 60 °C. Ultrathin sections (∼100 nm) were cut on a Reichert Ultracut E ultramicrotome, stained with 3% aqueous uranyl acetate for 25 min and Reynolds’ lead citrate for 5 min, and imaged using a FEI Tecnai transmission electron microscope at 80 kV.

### Airyscan imaging of actin cytoskeleton

Seeded cells on glass coverslips were fixed in 4% PFA at room temperature, washed three times with PBS, and stained with TRITC–phalloidin (Sigma-Aldrich P1951) for 20 min at room temperature. Samples were washed three times each with PBS and distilled water and mounted in Mowiol, then dried overnight. Actin was imaged on an LSM880 Airyscan confocal microscope in super-resolution mode using a 63× oil immersion objective (1.4 NA Plan-Apo), 540 nm excitation, 570 nm emission, and z-stacks acquired at 100 nm intervals over 10–20 µm.

Airyscan datasets were processed in Fiji. 3D projections were generated to visualise the dorsal actin network. Actin ridges and actin cortex were manually segmented and segmentations were validated by 3D re-projection. To minimise measurement artefacts introduced by ridge folding, extracted structures were reoriented to the yz plane prior to thickness quantification. Ridge detection was applied to reoriented images and visually inspected for its accuracy.

### Live cell imaging

#### Differential interference contrast

J774 cells (1 × 104 per well) were seeded in 96-well glass-bottom plates and incubated overnight at 37 °C, 5% CO_2_. Live imaging was performed on a Nikon Eclipse Ti widefield microscope in DIC mode using a 60× oil immersion objective (1.4 NA Plan-Apo). Polariser was adjusted to optimise ridge contrast. Time-lapse sequences were acquired for 15 min at 1 frame s^−1^ using an Andor Neo SCC-02546 camera. Ridge lifetime and thickness were quantified in Fiji using ridge detection-based analysis.

#### Lattice light-sheet data

Previously acquired lattice light-sheet datasets (Prof. Jennifer Stow, University of Queensland) were manually processed to separate ridges from dorsal ruffles (Condon et al., 2018). Preciously described ridge detection analysis was used to identify, track and measure individual ridges.

#### Phagocytosis assays

Two strains of *Cryptococcus neoformans* were used: H99 (serotype A) and CAP59. Frozen bead stocks were recovered from −80 °C, plated on YPD agar, and grown at 28 °C for 2–3 days. Colonies were re-streaked onto fresh YPD agar and grown for an additional 2–3 days. A loopful of culture was suspended in 2 ml YPD in a 5 ml tube and rotated at 20 rpm at 28 °C. Prior to infection, H99 cells were washed three times in PBS. CAP59 cells were washed in 0.1% PBS–Tween to reduce aggregation. Yeasts were labelled with calcofluor white (Sigma 18909) at 1:100 dilution for 20 min at room temperature. For opsonisation, H99 cells were incubated with monoclonal antibody 18B7 (1 µl per 1 × 10^6^ CFU) for 1 h at room temperature with 20 rpm rotation. Cells were washed, adjusted to 1 × 10^6^ CFU ml^−^1, and resuspended in serum-free medium. J774 macrophages (1 × 10^5^ cells per well) were seeded on uncoated glass coverslips in 24-well plates overnight at 37 °C, 5% CO_2_. For infections, media was replaced with 1 ml yeast suspension at an MOI of 1:10. Wells were kept at 37 °C, 5% CO_2_ for 2 hours prior to fixation in 4% PFA for 10 min at room temperature. For time-course experiments, wells were fixed at 0, 5, 10, 20, 30, 60 and 120 min post-infection. After fixation, samples were quenched with 50 mM NH_4_Cl for 5 min and permeabilised with 0.1% Triton X-100 for 3 min. Capsules were labelled by incubation with 18B7 for 20 min at room temperature, followed by anti-mouse IgG–FITC for 20 min and phalloidin–TRITC for 20 min at room temperature. Samples were washed with PBS and distilled water and mounted in Mowiol. Imaging was performed on a Nikon Eclipse Ti using a 60× oil immersion objective (1.4 NA Plan-Apo) in DAPI (excitation 436/20 nm, emission 480/40 nm), GFP (excitation 470/40 nm, emission 525/50 nm) and Cy3 (excitation 545/25 nm, emission 605/70 nm) channels..

To assay phagocytosis under conditions of ridge repression, J774 cells were first incubated with CAP59 for 1 h at MOI 1:10, then challenged with either H99, CAP59, or beads using the same MOI. Newly added H99 and CAP59 were prelabelled with calcofluor white to identify them from previously added cryptococcal cells. Controls included parallel infections with H99 or CAP59 without CAP59 pre-incubation. Samples were fixed after 2 h and processed for immunofluorescence and uptake scoring as described previously.

## Code availability

The ridge curvature profiling script is available at GitHub: https://github.com/jaimecanedo2025/ridgecurvature/blob/a9cbd6d210b2d7795c0de5a16506ac8e9b99afde/jamesCurvatureScriptv2.py

## Data availability

The contingency tables underlying attachment analyses are provided in Supplementary Tables S1–S4. Other data supporting the findings of this study are available from the corresponding author upon reasonable request.

## Statistics and reproducibility

Statistical analyses were performed using tests specified in the figure legends. Two-sided Fisher’s exact tests were used to compare proportions in contingency tables (Supplementary Tables S1–S4). Unpaired t-tests and multiple unpaired t-tests were used for comparisons between two groups as indicated (for example, phagocytic uptake assays). Ordinary one-way ANOVA was used for comparisons across three or more conditions. For non-normally distributed data, nonparametric tests were used as indicated (for example, Mann–Whitney). Exact sample sizes (n), definitions of n (cells, ridges, or independent experimental repeats), summary statistics (mean ± s.d. or median) and all statistical parameters (test statistic, degrees of freedom and P value) are reported in the figure legends.

## Ethics statement

Human samples were obtained in accordance with ethical practice approved by the **South Sheffield Research Ethics Committee**, with **written informed consent** from each donor. All animal experiments were performed in accordance with UK legislation under the Animals (Scientific Procedures) Act 1986. The University of Sheffield Project Review Committee approved all animal experiments which were carried out under the UK Home Office project licence P4802B8AC and UK Home Office Project Licence P5395C858.

## Acknowledgments

We would like to thank Chris Hill and Svet Tzokov of the University of Sheffield electron microscopy facility and Darren Robinson of the Wolfson Light Microscopy Facility for their support and training.

## Authors contributions

The project was conceptualised by SAJ, JC and RH, methodology was designed by SAJ, JC and JB, sample preparation and imaging were carried out by JC, MS, LMC, CN, YXH, data analysis was done by JC with critical advice from SAJ and RH, project supervision was carried out by SAJ and RH, project funding was secured by SAJ and RH, specimens were provided by SAJ, MS, JK, YXH, LMC, CN, LP, HW and EKT, original draft was written by JC, SAJ and RH before being reviewed and edited by all authors.

## Data, code, and materials availability

All data are available in the manuscript or the supplementary materials.

