## Supplemental data for "Macrophages use sub-micron actin ridges in phagocytosis of encapsulated fungal pathogens"

**Supplemental Figures**

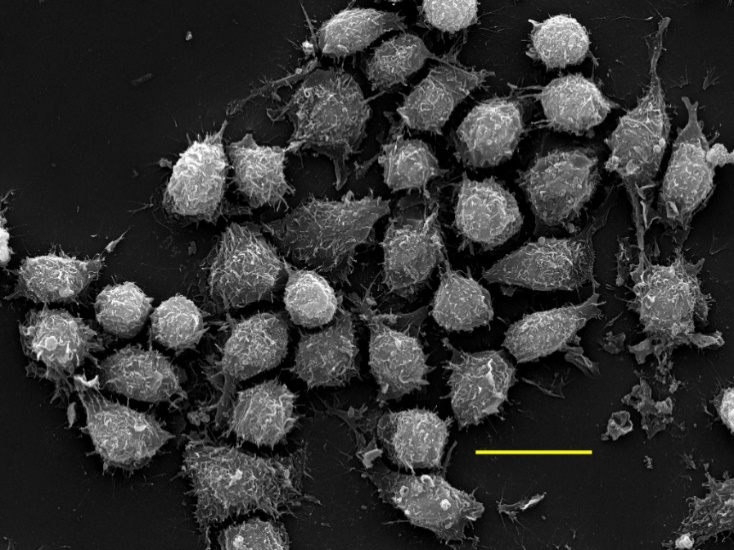

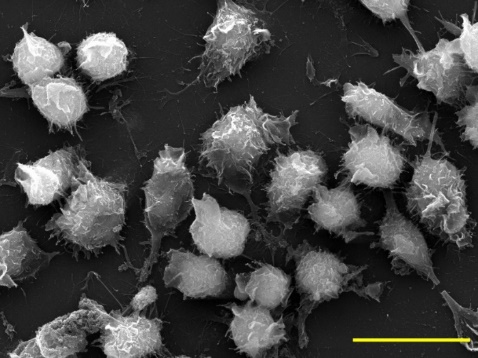

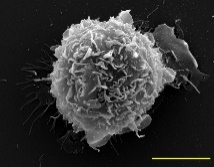

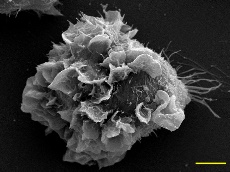

**B**

**A**

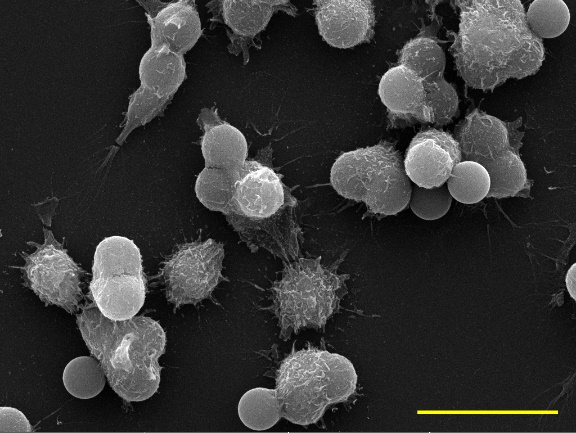

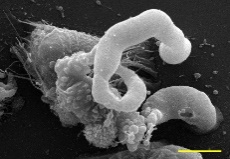

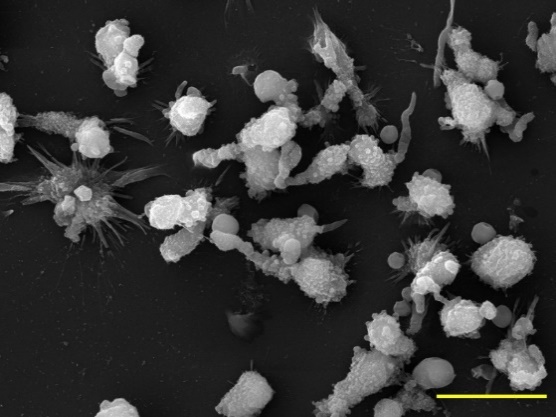

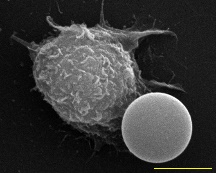

**C**

**D**

**F**

**E**

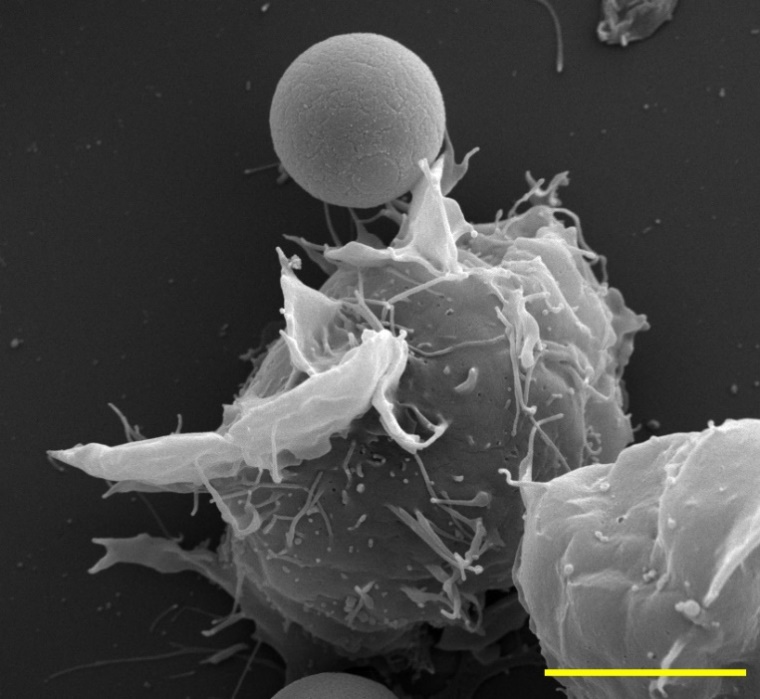

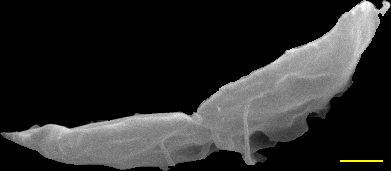

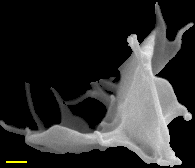

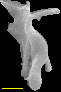

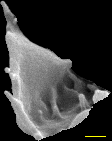

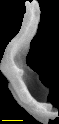

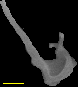

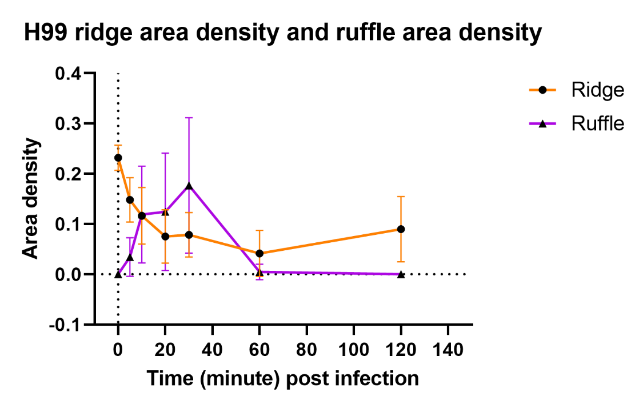

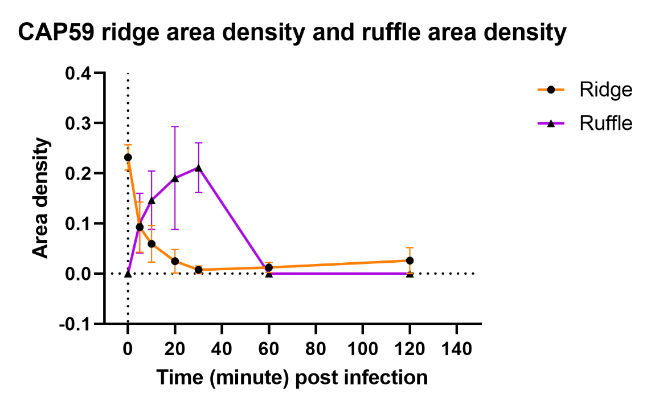

**H**

**G**

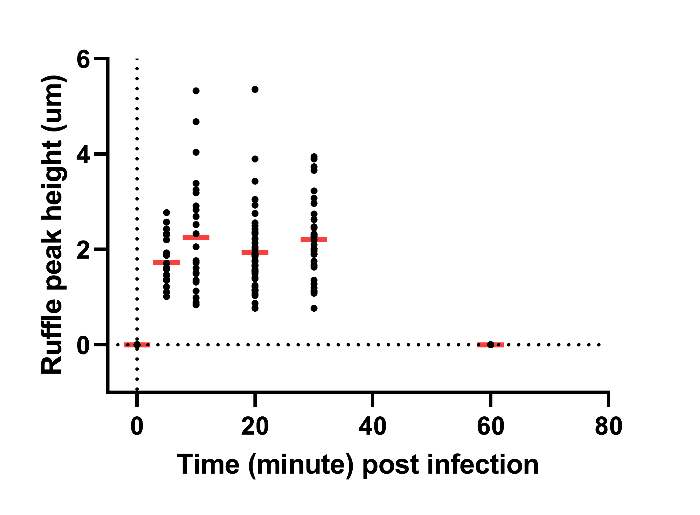

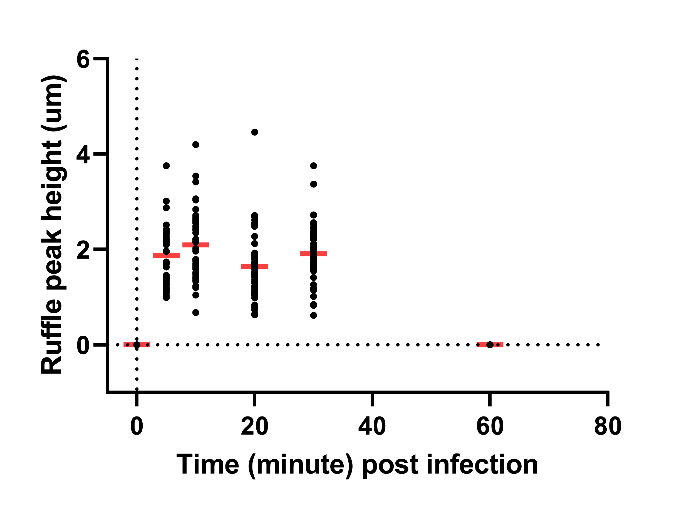

**I**

**J**

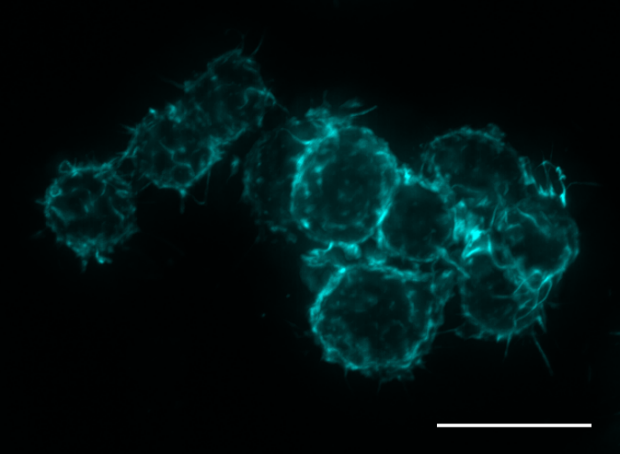

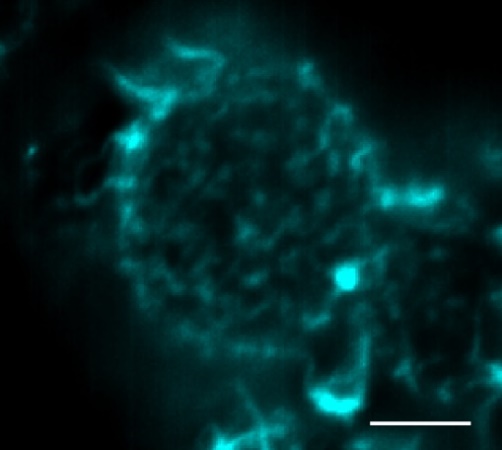

**K**

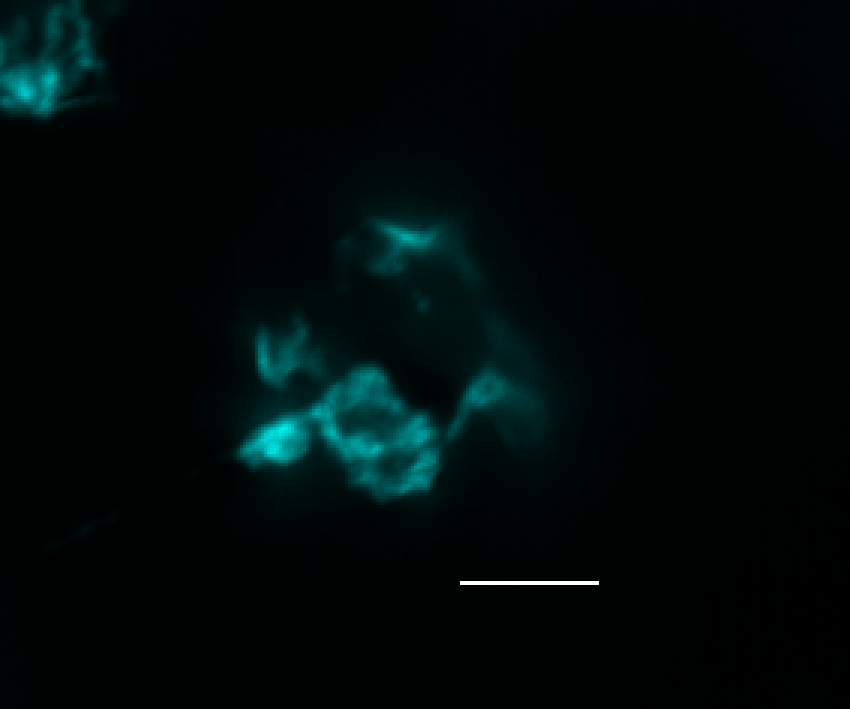

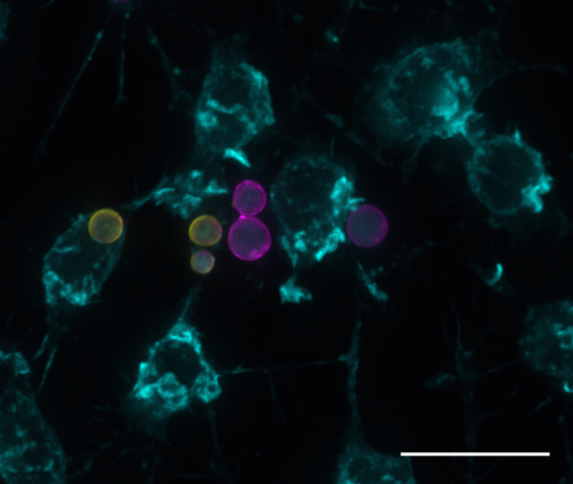

**L**

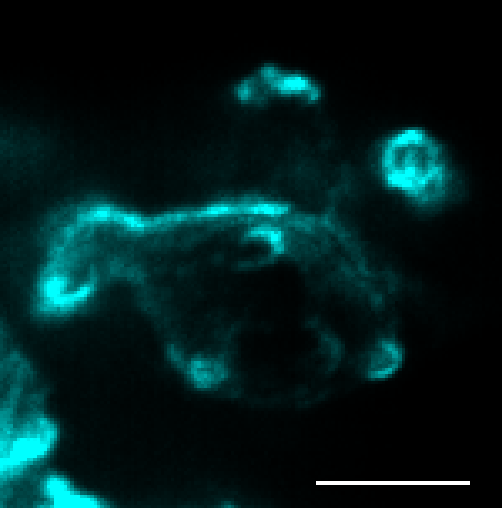

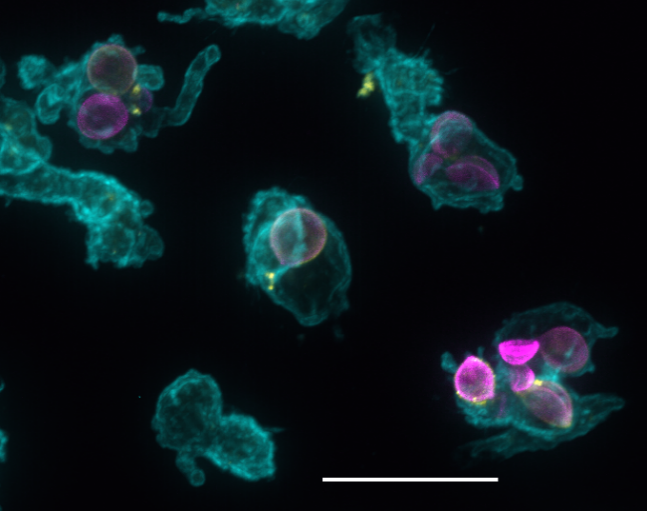

**M**

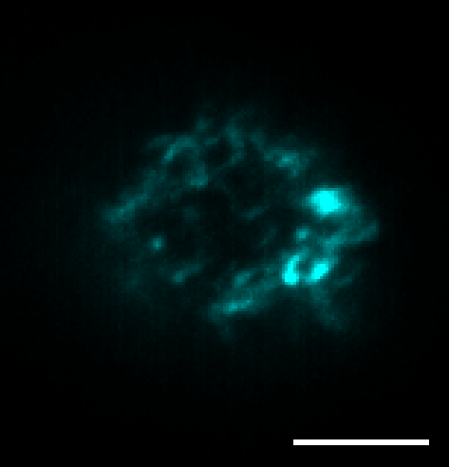

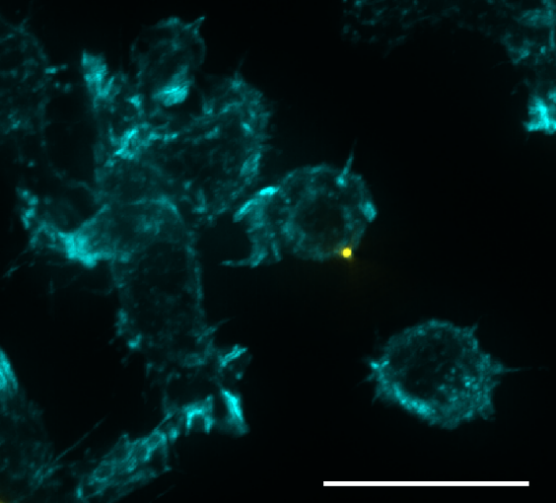

**N**

**Figure S1. Exposure of J774 macrophage to Cryptococcus reduces surface ridges while inducing membrane ruffles. a-c,** SEM images of J774 cell populations *in vitro* after incubating with *Cryptococcus neoformans* (H99) at different timepoints, and a representative cell is displayed in each inset to highlight the most common surface feature: **(a)** 0 minute post infection, the J774 surface has high density of ridges (inset arrow), **(b)** 30 minutes post infection, J774 surface is predominated by very distinct giant ruffles (inset arrow), and **(c)** 120 minutes post infection, a combination of bleb-like (inset yellow arrow) and tail-like (inset green arrow) surface features were observed in most cells, **d**, SEM image of J774 cells challenged with 6 um beads (inset yellow arrow) showing the persistence of ridges (inset green arrow) after 30 minutes, **e,** single cell SEM imaging using low accelerating voltage (10 kV) to resolve the nanoscopic surface structures of a J774 cell during its interaction with Cryptococcus (white arrow), **f,** manual extraction of ruffles produced by J774 cell in *d* to examine morphological properties of individual ruffles, the image border is colour-coded to easily identify which ruffle (arrow) it corresponds to in *d*, **g-h,** the production of membrane ridges and membrane ruffles in macrophages displayed inverse relationship in the first 30 minutes of infection: ridge density declined abruptly concomitant with a sharp increase in ruffle production in both wild type H99 **(g)** and acapsular CAP59 *C. neoformans* **(h)** infection models (*n* = 10-15 cells per timepoint, points: mean (*M*), bars: standard deviation), **i-j,** ruffle peak height was invariable throughout infection in both H99 **(i)** and CAP59 **(j)** infection models (*n* = 22-47 ruffles per timepoint, bars: *M*), **k-n,** cellular actin filament features of J774 cells using phalloidin-TRITC and widefield microscopy with blind deconvolution to examine actin during infection, adjacent image in yellow box shows the dorsalmost (1.5 um thickness) actin features of a representative cell (yellow box) from the main panel: **(k)** J774 cell actin cytoskeleton (cyan) challenged with H99 *C. neoformans* (capsule: yellow, cell wall: magenta) at 0 minute post infection, **(l)** 30 minutes post infection and (**m)** 120 minutes post infection, **(n)** J774 cell actin cytoskeleton (cyan) challenged with beads (yellow) for 30 minutes as negative control. Scale bars, 20 μm **(a-d** main panel**, k-n** main panel**)**; 5 μm **(a-d** inset**, k-n** inset**, e )**; 500 nm **(f)**.

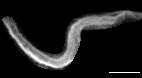

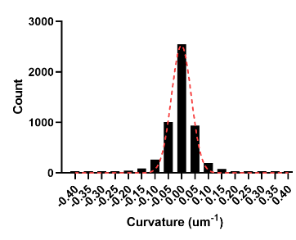

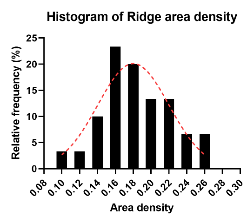

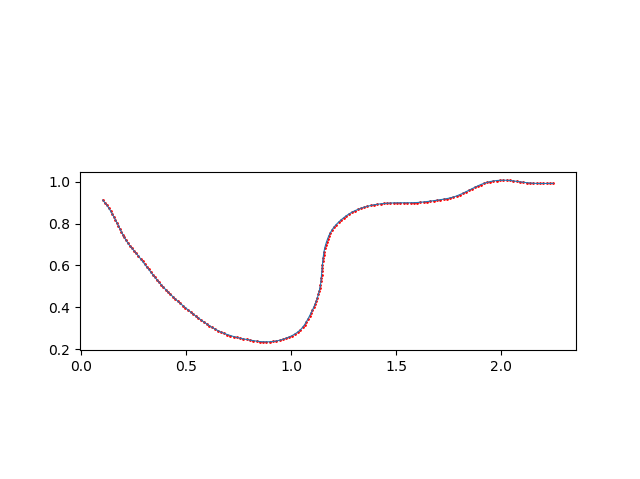

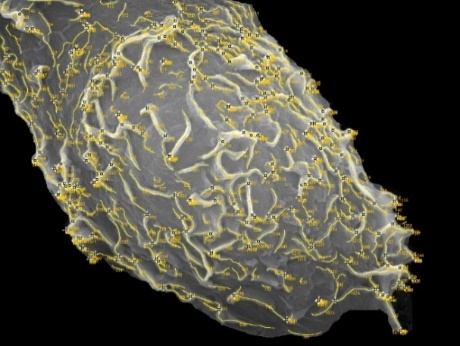

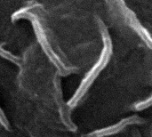

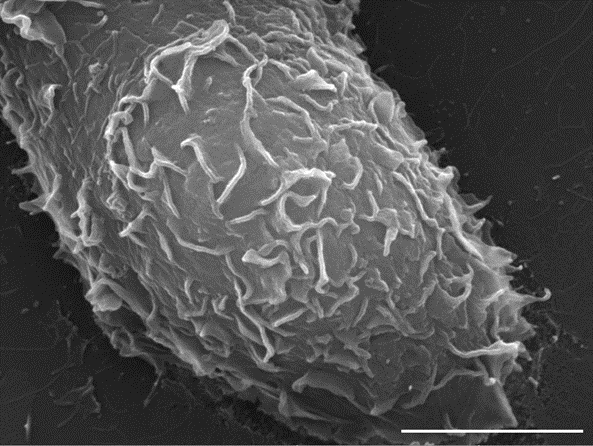

**B**

**A**

**D**

**C**

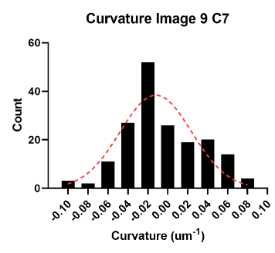

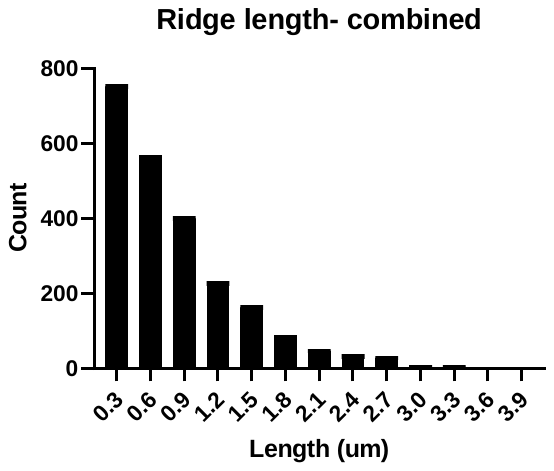

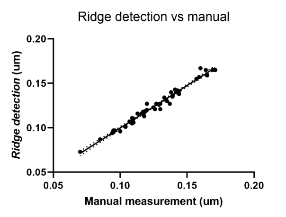

**I**

**H**

**G**

**F**

**E**

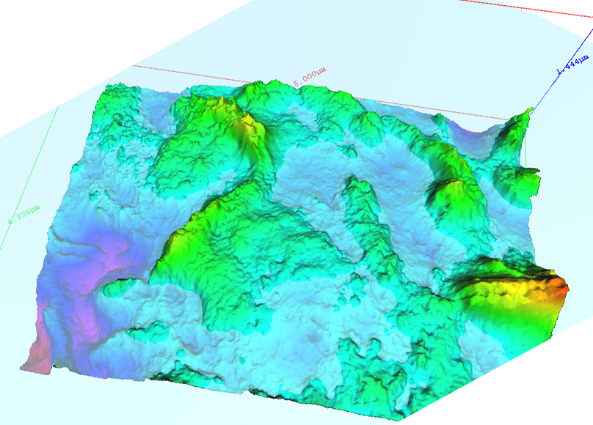

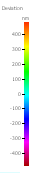

**J**

**K**

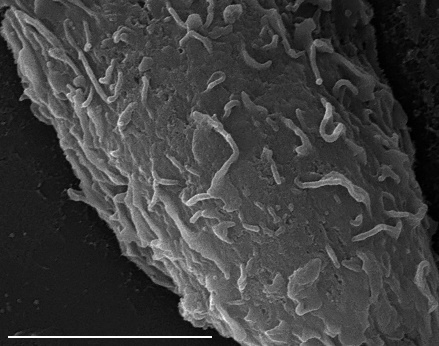

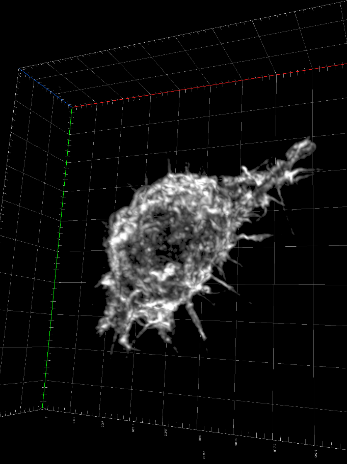

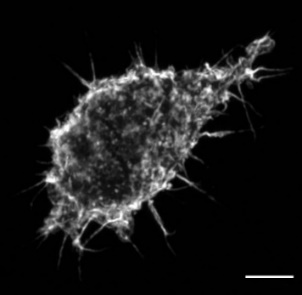

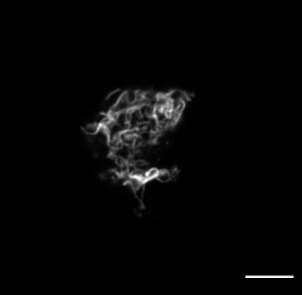

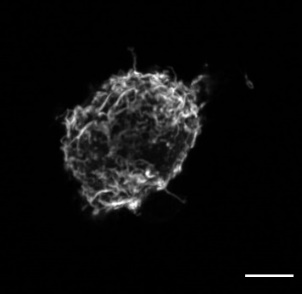

**L**

**M**

**N**

**O**

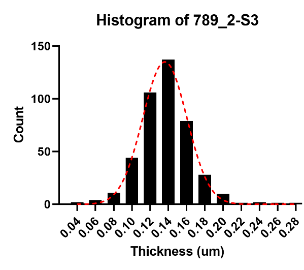

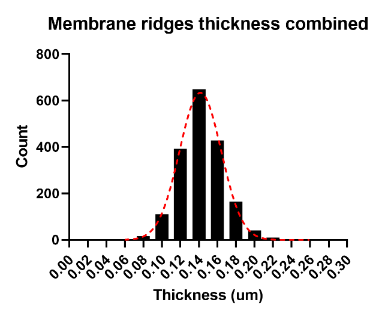

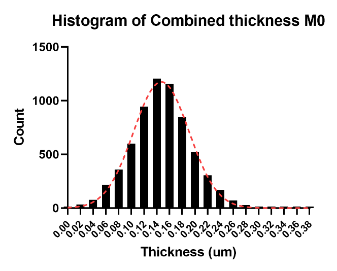

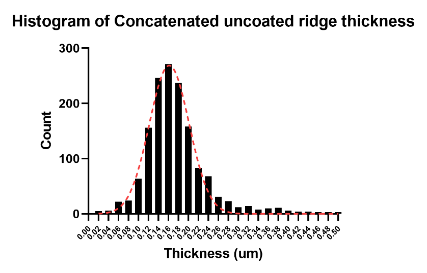

**P**

**R**

**S**

**Q**

(Figure legend on next page)

**Figure S2. Membrane ridges are dorsal sheet-like protrusions without any bias in curvature and with a highly conserved thickness across different phagocytes examined. a,** top view of a J774 macrophage surface revealing a high density of membrane ridges (inset: two membrane ridges with green line indicating thickness), **b,** image *a* processed with *ridge detection* (RD) plugin and showing ridges as numbered yellow lines, **c,** membrane ridge manually extracted from an SEM image, **d,** image *c* processed with RD and an improvised curvature profiling script to extract curvature properties, **e,** comparison of manual and automated (RD) measurements of membrane ridges revealing high measurement correlation (*R*^2^ = 0.978, *p* < 0.0001), **f,** combined ridge length profile in all SEM images of J774 cell (*n* = 12 cells (1818 ridges), mean (*M*) = 0.841, median = 0.670, skewness (*g_1_*) = 2.207), **g,** area density profile of membrane ridges in J774 cells with a Gaussian fit (*R*^2^ = 0.793, *n* = 30 cells, *M* = 0.18, *SD* = 0.04), **h,** ridge curvature profile of image *c* with Gaussian fit (*R*^2^ = 0.738) and showing slightly negative mean curvature (*M* = -0.01, *SD* = 0.04), **i,** combined curvature profile of all manually extracted ridges from SEM images of J774 cells (*R*^2^ = 0.989, *n* = 58 ridges (5220 ridge curvature points), *M* = 0.00, *SD* = 0.06), **j,** SEM image of a J774 cell surface used for photometric stereoscopy, **k,** 3D reconstruction of an area in *j* with a colour-coded depth map showing ridge thickness (red line) is conserved despite variable ridge height, **l,** maximum intensity projection of a J774 actin cytoskeleton imaged using phalloidin-rhodamine probe and confocal laser scanning microscope, **m-o,** three equal z section volumes of *l* each with about 1.5 μm thickness: **(m)** upper third/ dorsalmost section revealing actin ridges (green arrow), **(n)** middle section showing actin ridges (green), actin cortex (yellow) and actin filopodia (red), **(o)** lower third/ ventralmost section exhibiting more prominent actin cortex (yellow), actin filopodia (blue) and podosome-like structures (blue), **p-s,** thickness profiles of membrane ridges in different phagocytic cells with Gaussian fit (red curve): **(p)** J774 (*R*^2^ = 0.997, *n* = 12 cells (1818 ridges), *M*=142, *SD* = 24), **(q)** human monocyte-derived macrophages (*R*^2^ = 0.994, *n* = 9 cells (6555 ridges), *M*= 147, SD = 47), **(r)** mouse BMDM (*R*^2^ = 0.996, *n* = 4 cells (425 ridges), *M* = 137, *SD* = 28), **(s)** human neutrophils (*R*^2^ = 0.981, *n* = 11 cells (1467 ridges), *M* = 174, SD = 62). Scale bars, 5 μm **(a, j, m-o)**; 500 nm **(c).**

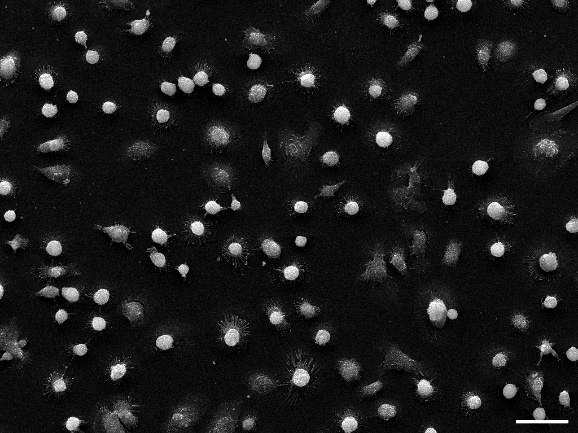

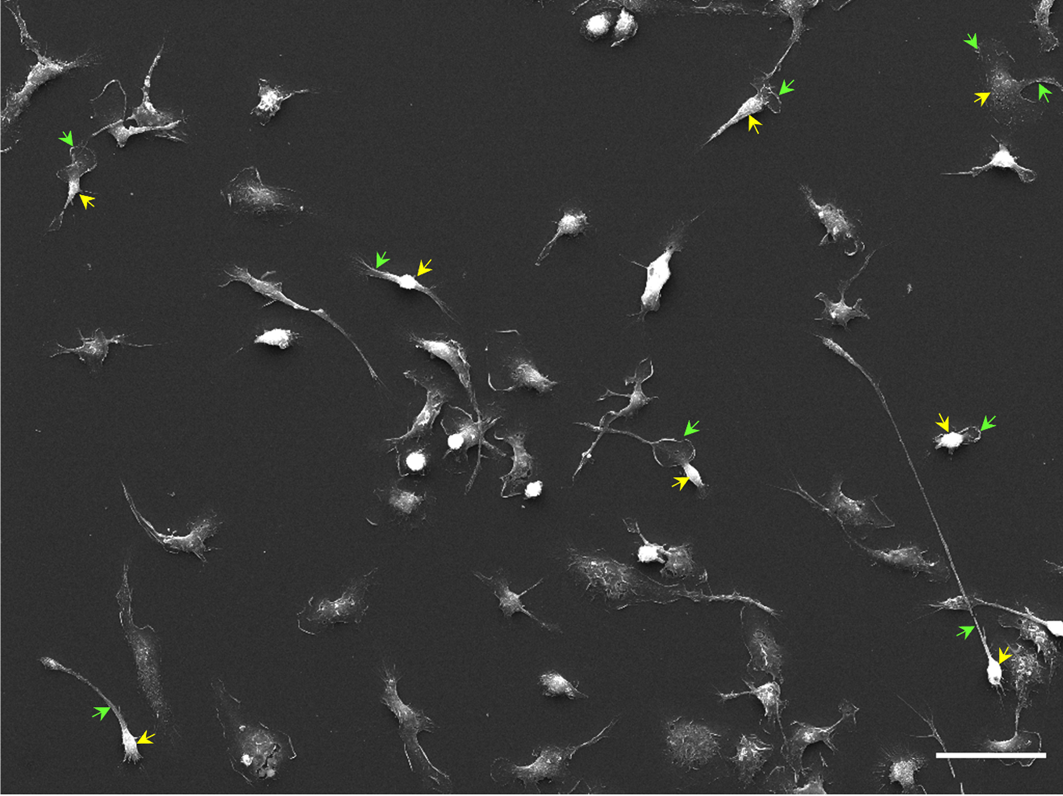

**B**

**A**

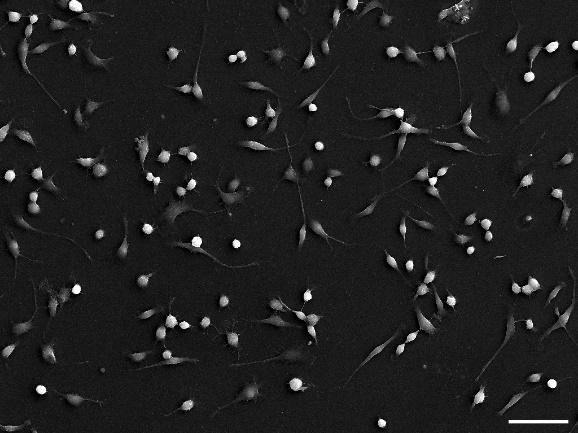

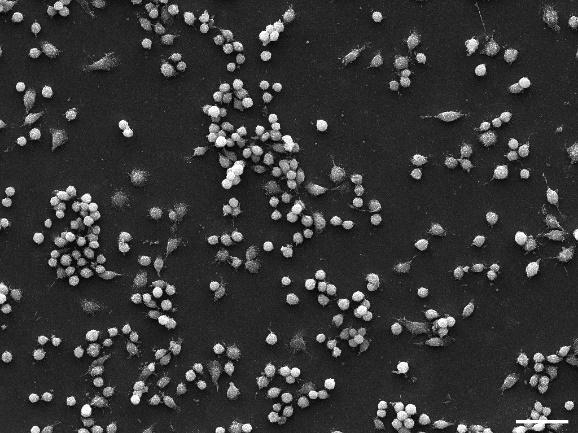

**D**

**C**

**Figure S3. Phagocytes assume different shapes and possess different types of membrane protrusion *in vitro*. a-d,** Low magnification SEM images revealing cellular morphologies of different phagocytes cultured *in vitro*: **(a)** Bone marrow-derived macrophages (BMDM) showing typical features of a phagocyte in SEM characterised by a distinctive bright cell body (yellow arrow) and a less bright membrane protrusion (green arrow), **(b)** human monocyte-derived macrophages (HMDM), **(c)** J774 macrophages, **(d)** RAW264.7 macrophages. All scale bars, 50 μm.

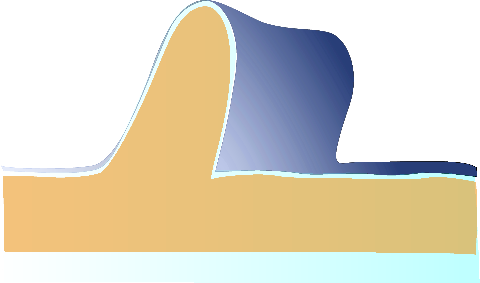

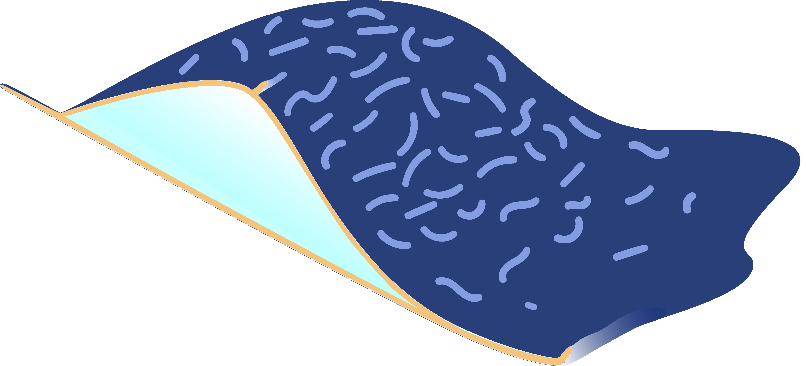

**B**

**A**

**D**

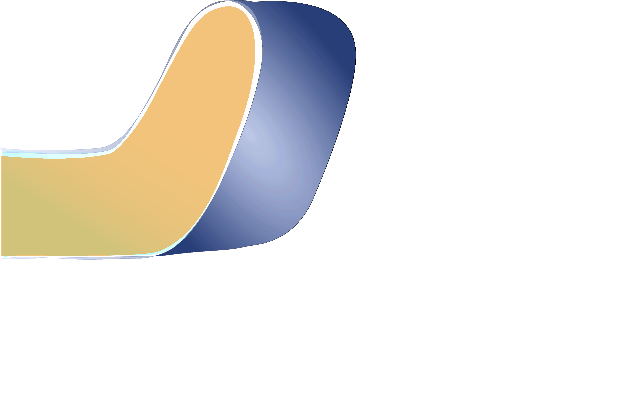

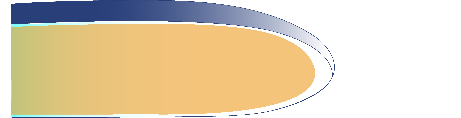

**C**

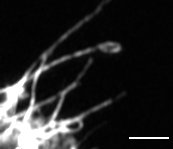

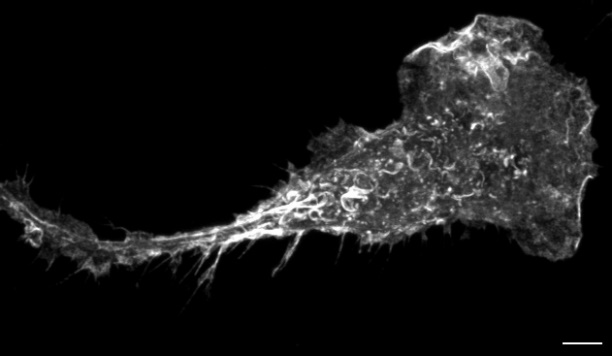

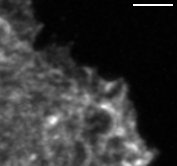

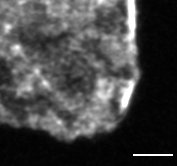

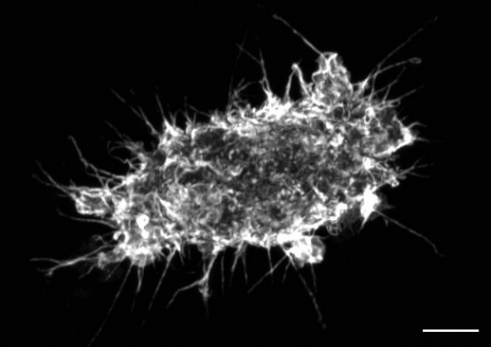

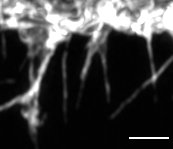

**F**

**E**

**G**

**H**

**I**

**J**

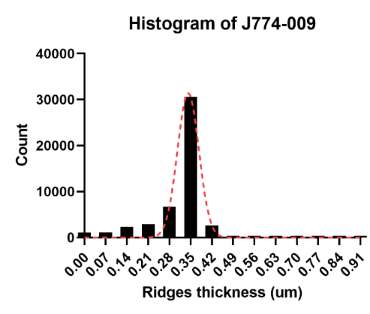

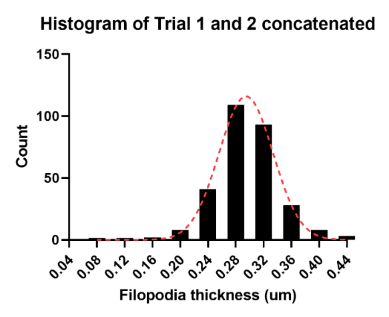

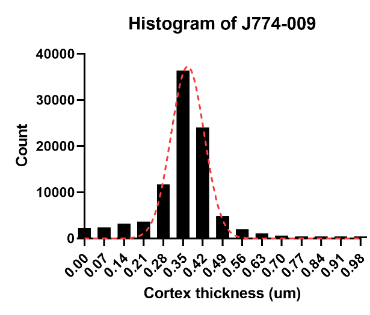

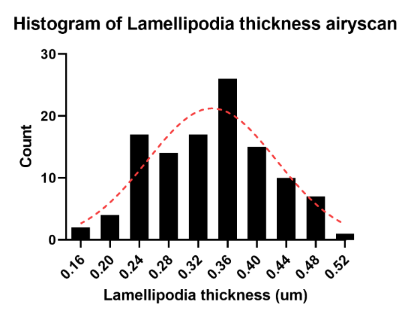

**L**

**M**

**K**

**N**

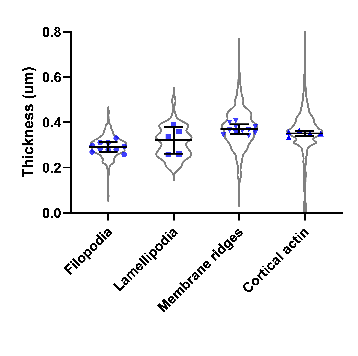

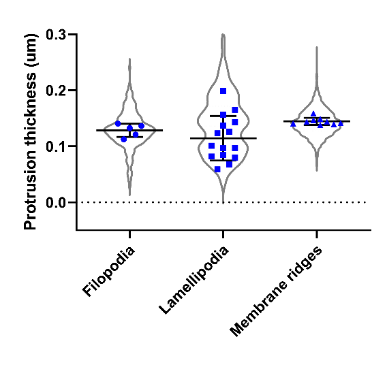

**O**

**P**

(Figure legend on next page)

**Figure S4. Surface ridges have similar thickness with filopodia and lamellipodia.** **a,** schematic diagram of a macrophage dorsoventral section showing its different types of actin-based membrane protrusion: **b,** zoomed-in diagram of a membrane ridge illustrating the thicknesses of its actin component (green double arrow) and the underlying actin cortex (red double arrow), **c,** zoomed-in diagram of a filopodium and its thickness (green double arrow), **d,** zoomed-in diagram of a lamellipodium with its thickness measured through local thickenings on its leading edge (figure 1a yellow arrow, figure s3g green arrow), **e,** maximum intensity projection of a J774 actin cytoskeleton imaged using airyscan superresolution laser scanning microscope and phalloidin-rhodamine probe, **f, g,** zoomed-in regions of the leading edge of *e* with occasional actin thickening regions (*g* green arrow), **h,** maximum intensity projection of a J774 actin cytoskeleton with numerous filopodia, **i, j,** zoomed-in regions of filopodia in *h*, **k-n,** thickness profiles of actin-based structures of J774 cells showing a Gaussian curve: **(k)** lamellipodial actin (*R*^2^ = 0.830, *n* = 5 cells, *M* = 312, *SD* = 66), **(l)** filopodial actin (*R*^2^ = 0.997, *n* = 10 cells, *M* = 294, *SD* = 46), **(m)** membrane ridge actin (*R*^2^ = 0.981, *n* = 11 cells, *M* = 369, *SD* = 86), **(n)** cortical actin (*R*^2^ = 0.978, *n* = 11 cells, *M* = 349, *SD* = 112.6), **o,** no statistically significant difference was found between thicknesses of different J774 cell protrusions using SEM images as confirmed by ordinary one-way ANOVA (*F*(2,26)=2.910, *p*=0.072), *n* = 5-15 cells, bars: *M* ± *SD*), **p**, no statistically significant difference was found between actin thicknesses using airyscan images of different J774 cell protrusions as confirmed by ordinary one-way ANOVA (*F*(2,23)=16.17, *p* =0.225), *n* = 5-11 cells, bars: *M* ± *SD)*,. Scale bars, 5 μm **(e, h)**; 2 μm **(f, g, i, j).**

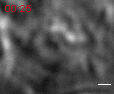

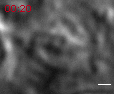

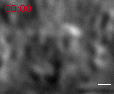

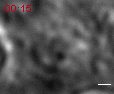

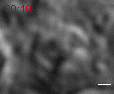

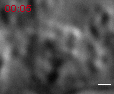

**C**

**B**

**A**

**F**

**E**

**D**

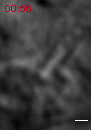

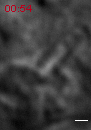

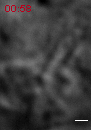

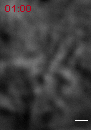

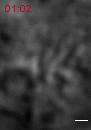

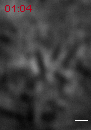

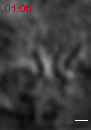

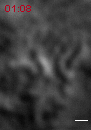

**J**

**I**

**H**

**G**

**N**

**M**

**L**

**K**

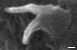

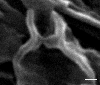

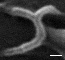

**Q**

**P**

**O**

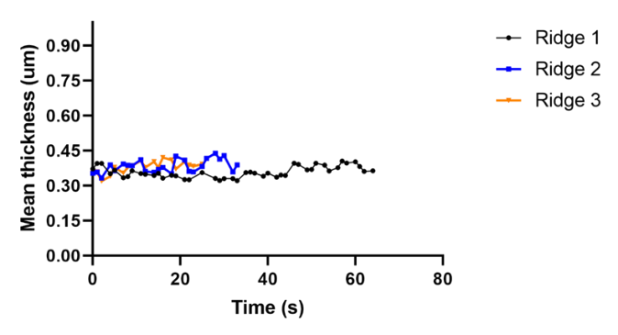

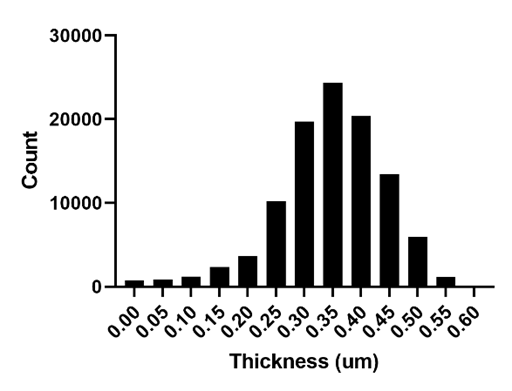

**S**

**R**

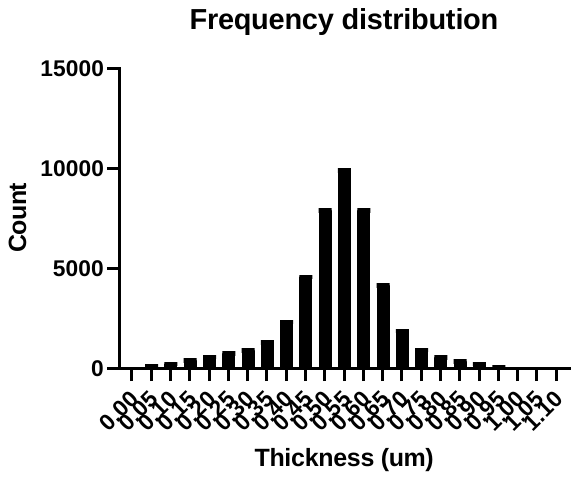

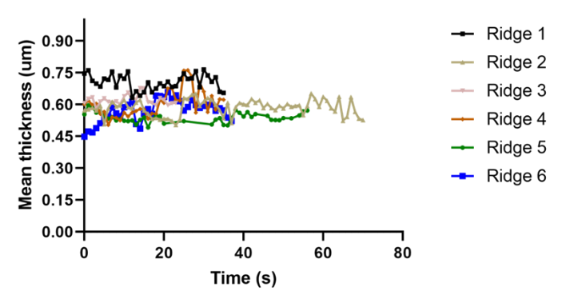

**U**

**T**

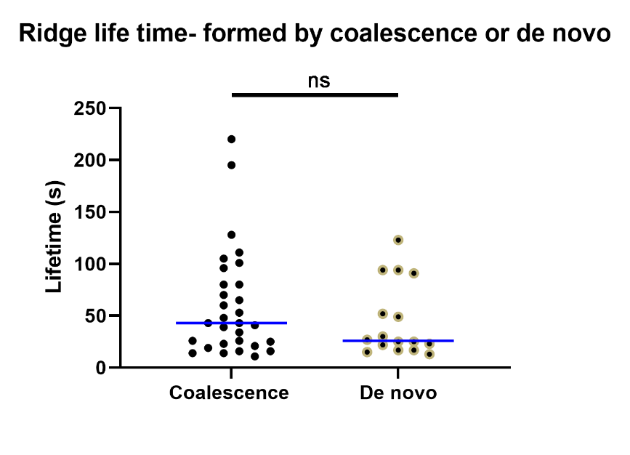

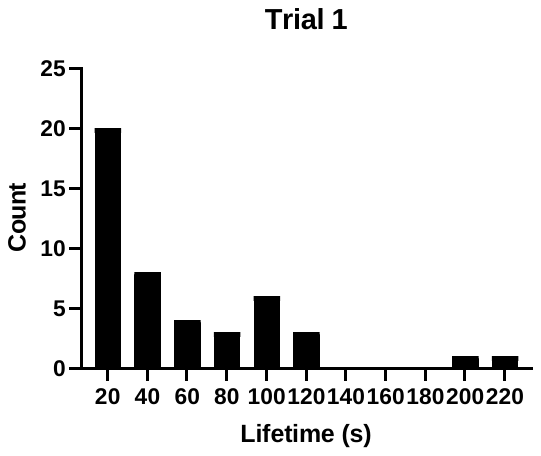

**W**

**V**

**Figure S5. Surface ridge is a dynamic structure capable of fusing and splitting, and its thickness is highly maintained throughout its lifetime. a-f,** DIC timelapse montage of membrane ridges with its lateral outgrowths (yellow arrow) which facilitate fusion before coalescing into a single ridge (green arrow) – see supporting video Movie 3 for complete timelapse data, **g-n,** timelapse montage of a surface ridge (yellow arrow) splitting from one end up to the opposite end through its midline (yellow broken line) to produce two daughter ridges (green arrow) – Movie 4 for complete timelapse data, **o-q,** high resolution SEM images of ridges fusing **(o)** revealing lateral outgrowths/fusion regions (yellow arrow), and ridge splitting **(p, q)** into daughter ridges (green arrow) with conserved thickness, **r,** thickness profile of membrane ridge actin in live macrophages expressing GFP-LifeAct imaged by lattice light sheet for 30 minutes (*n* = 3 cells, *M*= 346, *SD* = 94), **s,** individual thicknesses of ridges in *r* tracked throughout their lifetimes, **t,** thickness profile of membrane ridges using label-free and live imaging DIC for 15 minutes (*n* = 2 cells, *M* = 530, SD = 139), **u,** individual thicknesses of ridges in *t* throughout their lifetimes, **v,** lifetime histogram of membrane ridges using DIC imaging (*n* = 2 cells (46 ridges), *M* = 55, SD = 46), **w,** ridges formed by fusion or coalescence did not show any statistically significant difference in lifetime with those formed *de novo* as confirmed by Mann-Whitney test (*n* = 16-30 ridges, *p* = 0.333, bars: median). Scale bars, 1 μm **(a-n)**; 200 nm **(p-q)**.

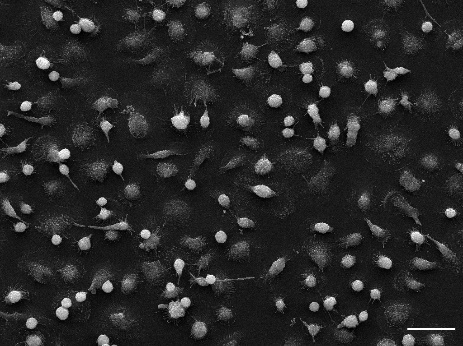

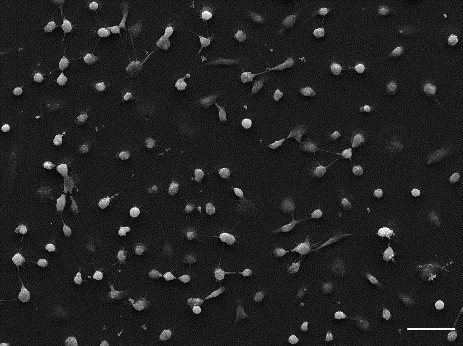

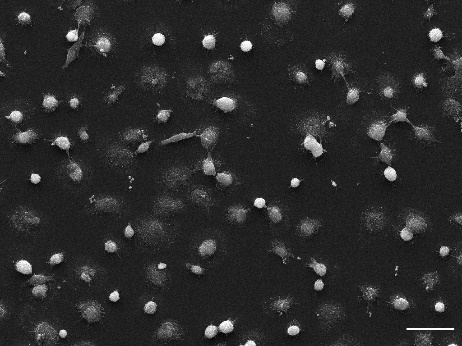

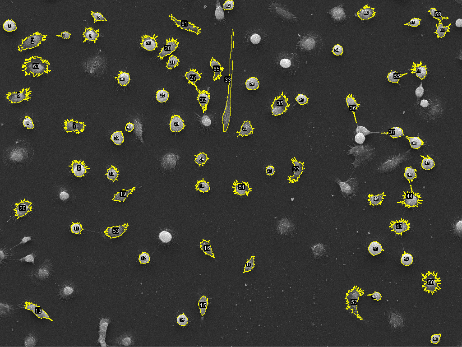

**A**

**B**

**D**

**C**

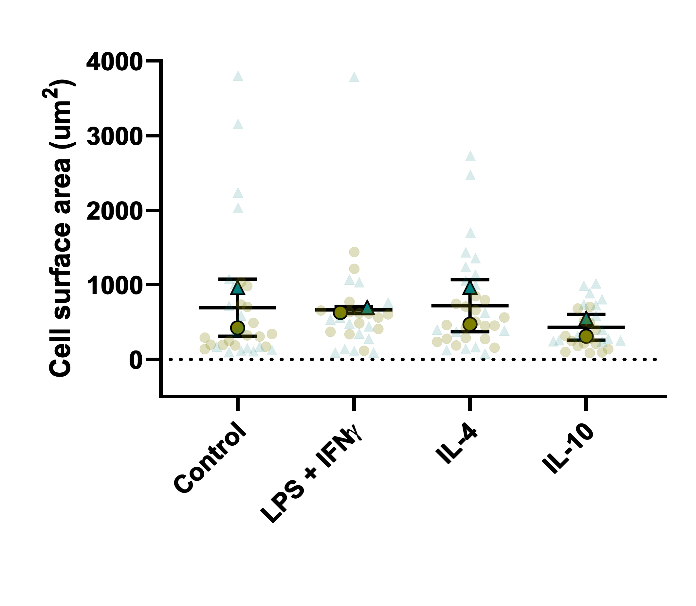

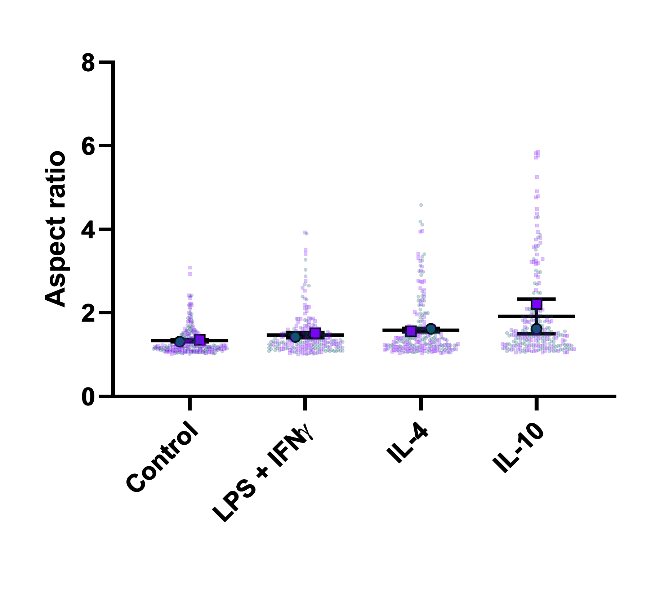

**F**

**E**

**Figure S6. No change in cell cross-sectional area nor cell aspect ratio (elongation) was observed after differentially activating human monocyte-derived macrophages (HMDM). a,** SEM image of unstimulated (control) HMDM *in vitro* showing cell borders (yellow outline) as determined by image segmentation using thresholding in Fiji, **b,** HMDM activated by LPS/IFNγ, **c,** by IL-4, and **d,** by IL-10, **e****,** no statistically significant difference in cell surface area of differentially-activated HMDM was found using ordinary one-way ANOVA (*F*(3,4) = 0.469, *p* = 0.72), *n* = 2 (14-17 cells per repeat), bars: mean (*M*) ± standard deviation (*SD*), **f,** no statistically significant change in aspect ratio of differentially-activated HMDM was found using ordinary one-way ANOVA (*F*(3,4) = 2.719, *p* = 0.179), *n* = 2 (96-140 cells per repeat), bars: *M* ± *SD*. Scale bar in all images, 50 μm.

**Tables**

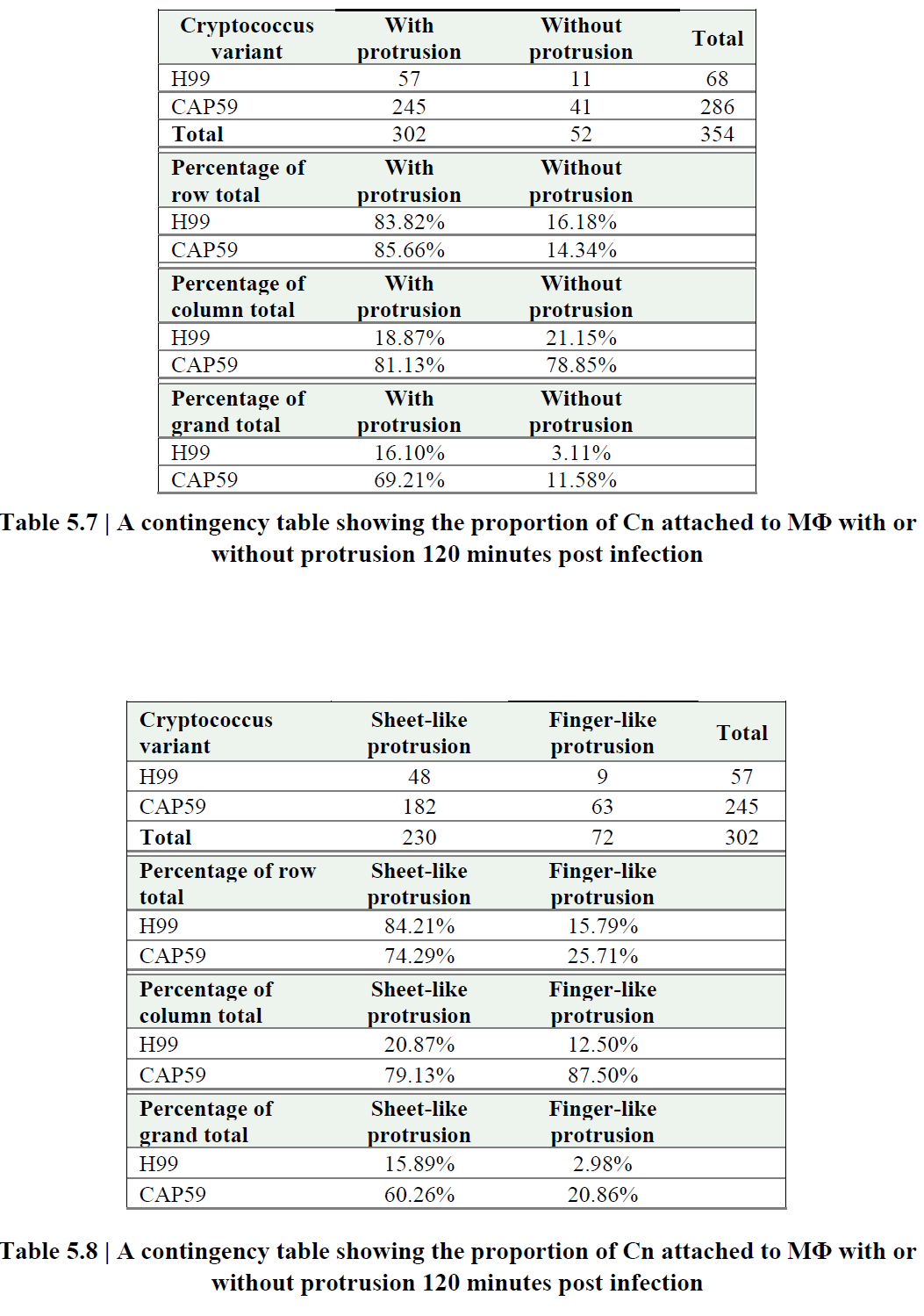

Table S1. A contingency table showing the proportion of capsular (H99) and acapsular (CAP59) Cryptococcus neoformans (Cn) attached to macrophages MΦ with or without protrusion after *in vitro* infection assay

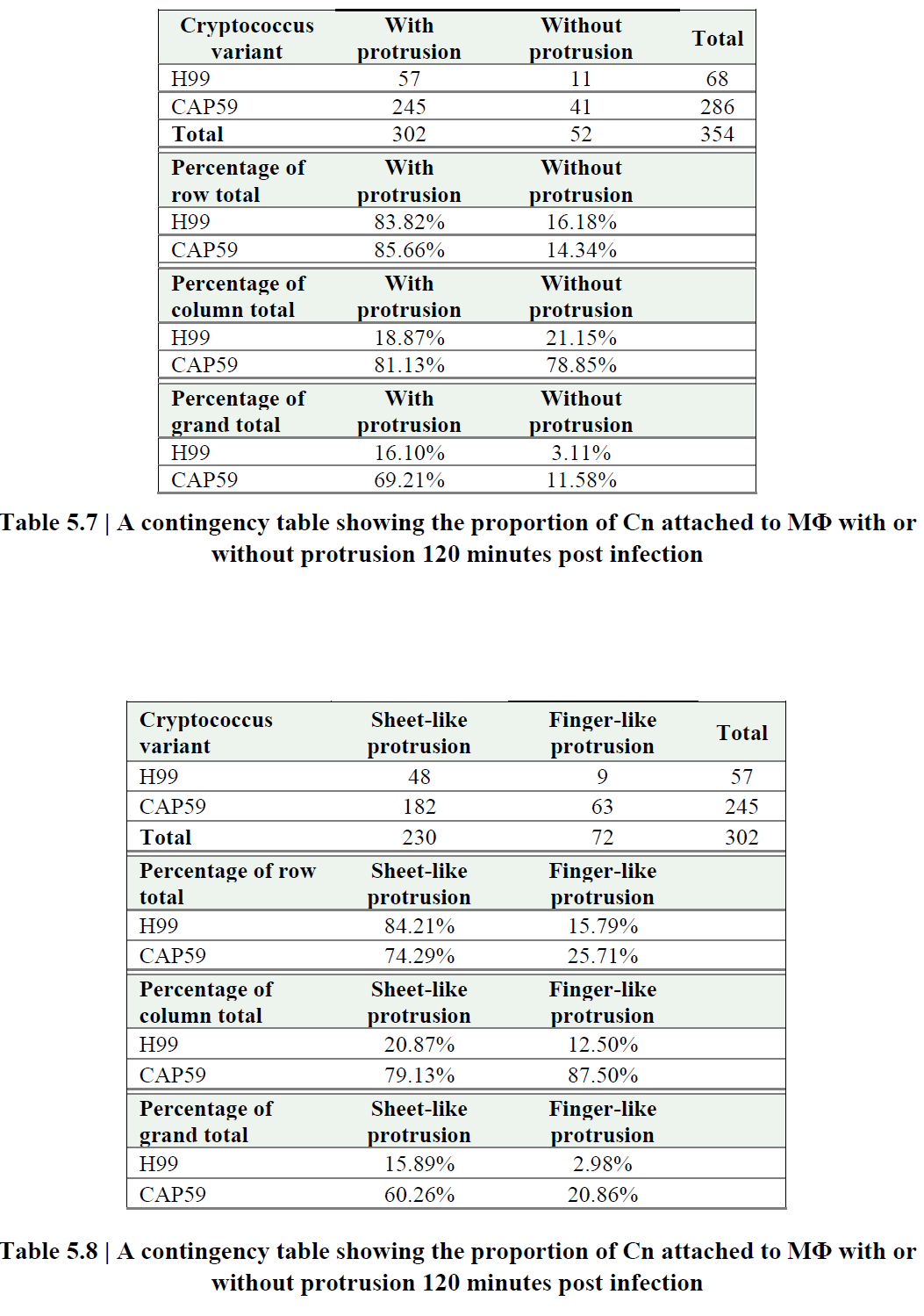

Table S2. A contingency table showing the proportion of capsular (H99) and acapsular (CAP59) Cryptococcus neoformans (Cn) attached to macrophages MΦ through sheet-like or finger-like protrusion after *in vitro* infection assay

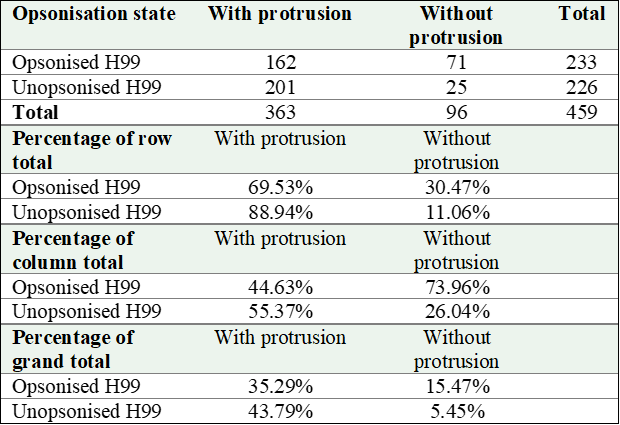

Table S3. A contingency table showing the proportion of opsonised or unopsonised H99 Cn attached to macrophages MΦ with or without protrusion after *in vitro* infection assay

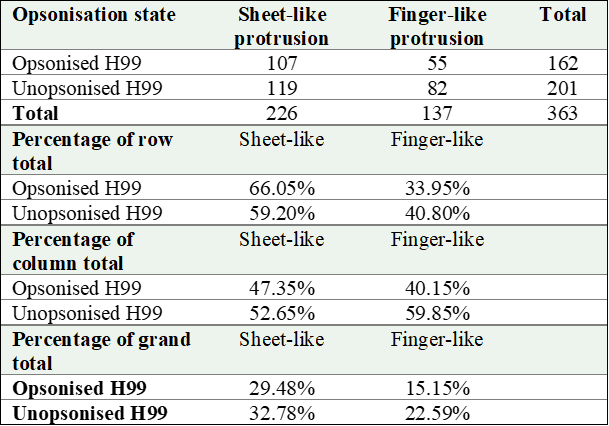

Table S4. A contingency table showing the proportion of opsonised or unopsonised H99 Cn attached to macrophages MΦ through sheet-like or finger-like protrusion after *in vitro* infection assay

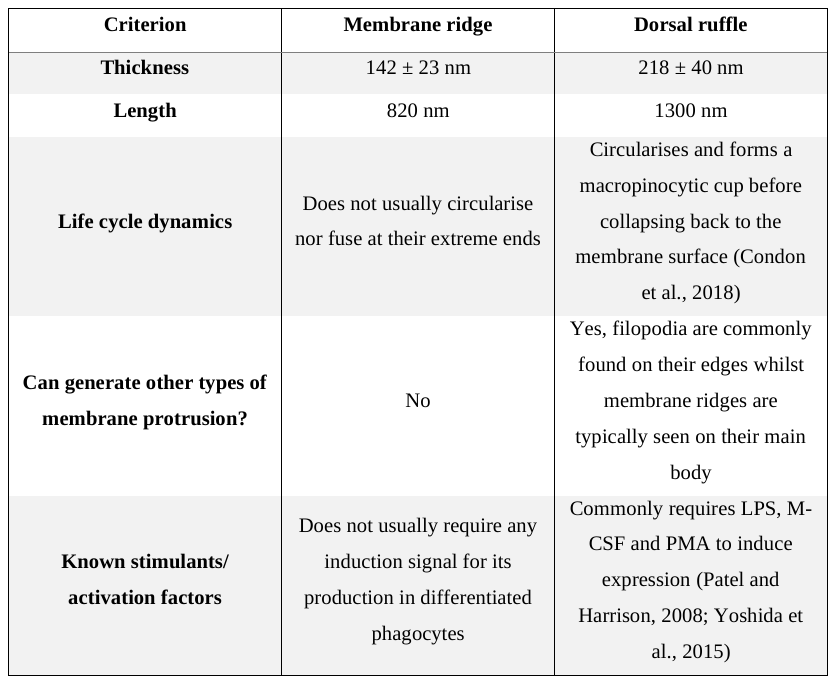

Table S5. Comparing membrane ridge and dorsal ruffle
